# Shared and Unique Evolutionary Trajectories to Ciprofloxacin Resistance in Gram-negative Bacterial Pathogens

**DOI:** 10.1101/2021.04.06.438548

**Authors:** Jaime E. Zlamal, Semen A. Leyn, Mallika Iyer, Marinela L. Elane, Nicholas A. Wong, James W. Wamsley, Maarten Vercruysse, Fernando Garcia-Alcalde, Andrei L. Osterman

## Abstract

The resistance to broad-spectrum antibiotic ciprofloxacin is detected in high rates for a wide range of bacterial pathogens. To investigate dynamics of ciprofloxacin resistance development we proposed a comparative resistomics workflow for three clinically relevant species of Gram-negative bacteria: *Escherichia coli*, *Acinetobacter baumannii*, and *Pseudomonas aeruginosa*. We combined experimental evolution in a morbidostat with deep sequencing of evolving bacterial populations in time series that reveals both shared and unique aspects of evolutionary trajectories patterns. Representative clone characterization by sequencing and MIC measurements enabled direct assessment of mutations impact on the extent of acquired drug resistance. In all three species we observed a two-stage evolution: (1) early ciprofloxacin resistance reaching 4-16-fold of wildtype MIC commonly as a result of single mutations in DNA gyrase target genes (*gyrA* or *gyrB*) and (2) additional genetic alterations affecting transcriptional control of drug efflux machinery or secondary target genes (DNA topoisomerase *parC* or *parE*).

**Importance:** The challenge of spreading antibiotic resistance calls for systematic efforts to develop more “irresistible” drugs based on deeper understanding of dynamics and mechanisms of antibiotic resistance acquisition. To address this challenge, we have established a comparative resistomics approach which combines experimental evolution in a continuous culturing device, the morbidostat, with ultradeep sequencing of evolving microbial populations to identify evolutionary trajectories (mutations and genome rearrangements) leading to antibiotic resistance over a range of target pathogens. Here we report the comparative resistomics study of three Gram-negative bacteria (*Escherichia coli*, *Acinetobacter baumannii*, and *Pseudomonas aeruginosa),* which revealed shared and species-specific aspects of the evolutionary landscape leading to robust resistance against the clinically important antibiotic ciprofloxacin. In addition to specific findings, the impact of this study is in highlighting the anticipated utility of a morbidostat-based comparative genomic approach to guide rational optimization of treatment regimens for current antibiotics and development of novel antibiotics with minimized resistance propensities.

## Introduction

Increasing antibiotic resistance is a premier threat to modern medicine. This danger necessitates expanded research into the mechanisms by which organisms gain resistance and continued development of new drugs to replace those becoming ineffective. A second-generation fluoroquinolone antibiotic, ciprofloxacin (CIP) was introduced for medical use in 1987 and boasted a wider spectrum of efficacy than first-generation quinolones (1). Ciprofloxacin is commonly prescribed as a front-line treatment against a broad range of bacterial infections (2, 3). Quinolone antibiotics target bacterial DNA gyrase (GyrA/GyrB) and topoisomerase IV (ParC/ParE), enzymes which control DNA supercoiling during replication and transcription. By binding to these enzymes when they are complexed with DNA, CIP inhibits the repair of DNA breaks and causes irreversible damage to the genome (4–8).

A weak to moderate resistance to CIP can occur through a single missense mutation in one of the target genes yielding resistant variants, even at drug levels substantially below MIC (Minimum Inhibitory Concentration). Thus, a reported most common GyrA:S83L CIP-resistant variant of *E. coli* has emerged at the drug concentration of only 1/230 of MIC (9). Further stepwise acquisition of additional mutations in the presence of greater drug challenge (10, 11) confers increased resistance to fluoroquinolones (12, 13). The resulting highly resistant forms typically contain a combination of mutations in target genes (GyrAB and/or ParCE), species-specific efflux pumps (such as AcrABC in *E. coli*), and porins mediating drug influx (14–18).

The observed similarity of intrinsic CIP-resistance mechanisms in divergent target pathogens is consistent with the universal mechanism of action of this broad-spectrum drug. Nevertheless, distinct species display different resistibility potential with respect to the dynamics and extent of acquired resistance. The primary objectives of this study were to assess and compare the dynamics of CIP-resistance acquisition and resistance mechanisms in three divergent species representing difficult-to-treat Gram-negative bacteria, *Escherichia coli*, *Acinetobacter baumannii*, and *Pseudomonas aeruginosa* in a standardized setting of a continuous culturing device. Among these groups of pathogens, *A. baumannii* is of particular concern due to its genomic plasticity, a feature which gives rise to diverse isolates displaying preexisting and readily-acquired multiple drug resistance (19, 20). Another common nosocomial pathogen, *P. aeruginosa* is also known to cause dangerous antibiotic-resistant infections. In our comparative study, we have included the best studied model Gram-negative bacterium, *E. coli* K-12, a close relative of clinically relevant pathogenic strains of *E. coli*.

Although some mutations conferring CIP-resistance observed in clinical isolates and laboratory studies of all three target species were previously reported (10, 21, 22), a direct comparison of their evolutionary trajectories is complicated due to different selection conditions. In traditional experimental evolution studies, selection proceeds through a series of bottlenecks reflecting specifics of the experimental setup and bacterial population size. Thus, a serial transfer of small-size bacterial cultures leads to the predominant propagation of high-frequency/low-fitness mutants, while studies of large bacterial populations tend to yield low-frequency/high-fitness mutants that are commonly isolated from patients with CIP-resistant infections (10, 23). Another variable parameter defining the outcome of experimental evolution is the Mutant Selection Window (MSW)(24, 25). The MSW represents a drug concentration range enabling effective elimination of less resistant cells and propagation of more resistant cells. A morbidostat (constant morbidity) approach used in this study is based on an automated dynamic adjustment of the MSW, shifting upward over the course of experimental evolution (26, 27). This technique allows us to alleviate selection bottlenecks and enrich evolving bacterial populations with more robust resistant variants.

For a comparative CIP-resistomics analysis of the three selected Gram-negative bacteria in a standardized morbidostat-driven setup (**Figure 1a,b**), we leveraged an experimental evolution workflow developed and validated in our previous model study on evolution of triclosan resistance in *E. coli* (28) Briefly, the workflow used in this study **(Supplementary Figure S1)** includes: (i) competitive outgrowth of six parallel bacterial cultures in a custom-engineered continuous culturing device, the morbidostat, under gradually increasing antibiotic concentration; followed by (ii) sequencing of total genomic DNA from bacterial population samples taken as time series; (iii) identification and quantitation of sequence variants (mutations, small indels, mobile elements insertions) reflecting evolutionary dynamics and inferred resistance mechanisms; and (iv) experimental characterization of genotype-to-phenotype associations via mapping of mutations and determination of MIC values in selected individual clones.

**Figure 1.**
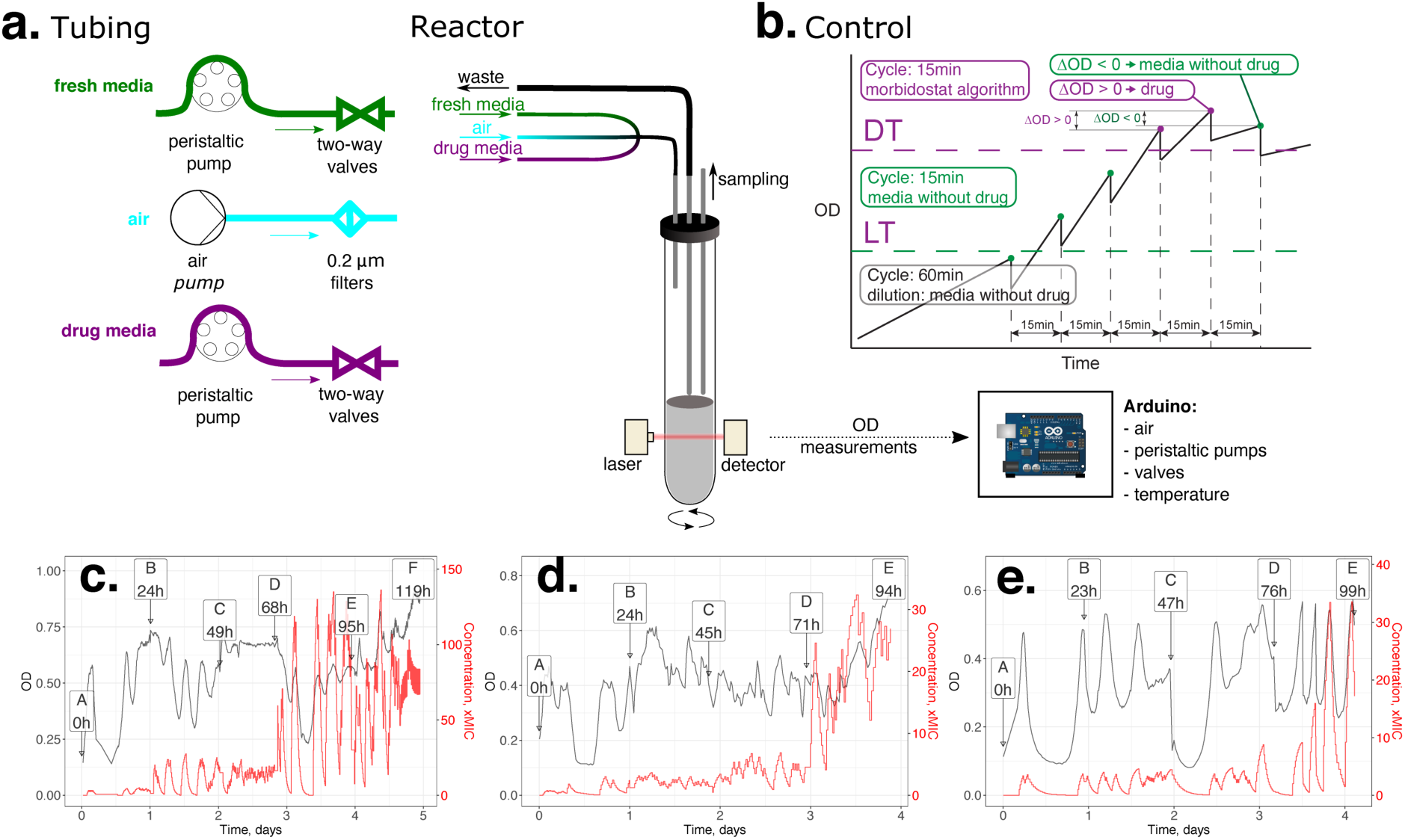
**Morbidostat design (a), control logics (b) and examples of evolutionary runs of *E. coli* (c)*, A. baumannii* (d) and *P. aeruginosa* (e) with ciprofloxacin. a.** Bacterial populations are continuously cultured in a 20 mL glass tube (bioreactor) with magnetic stirring and three input lines: filtered air (blue) and media from two feed bottles, with and without a concentrated drug (red and green, respectively). The growth (turbidity) is monitored using a laser beam and diode light sensor. Upon periodic addition of 2-4 mL media from the first or the second feed bottle (as defined by control logic, see B), the excess volume is displaced by air flow into a waste bottle. Samples (up to 10 mL) are taken periodically (1-2 times per day) through a dedicated sampling port. Our current morbidostat implementation includes 6 parallel bioreactors with individual feed lines that are independently monitored and controlled by the Arduino board with Windows PC-based user interface. **b.** Morbidostat logic is controlled by an Arduino board based on the principles described by Toprak et al. (27) using the real-time OD input from each bioreactor and predefined run parameters: lower threshold (LT), drug threshold (DT), and cycle time (time between dilutions, typically 10 - 20 min when OD>LT). Depending on the conditions shown in the diagram, one of the two peristaltic pumps (feeding media with or without drug) are engaged at the beginning of each dilution-outgrowth cycle. **c-e.** Representative OD profiles (black line) and calculated drug concentration profiles (red line) observed in the course of experimental evolution of *E. coli*, *A. baumannii*, and *P. aeruginosa* toward resistance against ciprofloxacin (CIP). One of the reactors is shown for each organism, while evolutionary profiles for all other experiments and reactors are provided in Supplementary Figure 2. The right axis shows CIP concentration (xMIC) as fold-change relative to MIC value of respective unevolved strains.

The performed analysis yielded the identification of mechanistically and clinically relevant CIP-resistance conferring mutations. This investigation revealed both shared and species-specific aspects of the evolutionary dynamics of resistance acquisition. It also confirmed the utility of the established workflow for comparative resistomics of known antibiotics, and potentially novel drug candidates, over a broad range of bacterial pathogens.

## Methods

### Morbidostat setup

Experimental evolution of CIP*-*resistance was performed using an optimized version of a morbidostat device, which was engineered and validated in our previous study on the evolution of triclosan resistance in *E. coli* (28). The general design is based on the principles described in (26, 27), extending the chemostat approach toward evolution of drug resistance. In the morbidostat, culture densities are maintained by regular automated dilutions with media containing or not containing antibiotic. This leads to gradual adaptation of bacterial populations to higher drug concentrations.

The detailed technical description of morbidostat hardware and accompanying software is provided in GitHub (https://github.com/sleyn/morbidostat_construction). Briefly, the main components of the device (**Figure 1a)** are: (i) six 20 mL glass tubes used as bioreactors, with magnetic stir bars and cap assemblies containing three air-tight needle ports for: (a) introduction of fresh media and continuous air flow for culture aeration, (b) liquid displacement after dilution, and (c) sample collection; (ii) silicone rings used to secure reactors in 3D-printed plastic housings which each contain a laser and a sensor diode for measuring culture turbidity; (iii) a small air-pump to provide aeration for growing cultures (fitted with 0.22 µm pore filters to block contamination in air feedlines) and enable liquid displacement from reactors (over a fixed level corresponding to a total volume of 20 mL); (iv) a thermoregulated heater and fan to control the temperature inside the morbidostat enclosure; (v) two 2 L bottles with tubing connecting each to a peristaltic pump which controls the flow of media (with and without antibiotic) to the reactors, where an assembly of 12 check valves (2 valves per reactor, each connected with one of the two pumps) controls delivery of media to individual reactors during each dilution; and (vi) a six-position magnetic stir plate which agitates cultures and enables mixing of media upon dilution. An Arduino-based microcontroller is programmed to control the following main parameters of the run: (i) enclosure temperature; (ii) time between dilutions (cycle time, CT); and (iii) selection of the volume delivered by Pump 1 (media without drug) or Pump 2 (media with drug) for each culture dilution, depending on the culture turbidity and growth rate in each tube. A user interface for parameter manipulation and real-time status display (including growth curves) is run on a PC using MegunoLink software (https://www.megunolink.com/, v. 1.17.17239.0827 for CEC-2, CEC-4 and CAB runs and v. 1.32.20005.0105 for PAC-1 and PAC-2 runs). Automated dilutions are controlled using the following encoded logic (**Figure 1b**):

1. In the active “dilution mode”, the optical density (OD) at the end of the current cycle (OD1) is compared with three parameters: (i) the predefined Lower Threshold (LT, typically ≤ 0.15); (ii) predefined Drug Threshold (DT, typically ∼ 0.3); and (iii) OD reached during the previous cycle (OD0). All OD values are corrected by adding OD of fresh media with antifoam.
2. The dilutions are always made by a predefined volume (typically V = 2 or 4 mL); if OD1 ≥ DT and (OD1-OD0) ≥ 0 the drug-containing media (Pump 2) is used with the interval corresponding to the predefined CT (typically 15–20 min); else drug-free media (Pump 1) is added.
3. If OD1 < LT, the system performs hourly dilutions with drug-free media. In the beginning of the run, this allows all six cultures to reach the same minimal density (OD1 = LT) prior to entering the active dilution mode. During the run, this “safe mode” prevents a complete wash-out of the culture after an excessive dose of drug.

### Morbidostat runs

Drug-free base medium consisted of Cation-Adjusted Mueller Hinton Broth (MHB)(Teknova) with a final concentration of 2% DMSO (except for CEC-2 and PAC-1 runs, which did not have DMSO) and 1/2,500 dilution of Antifoam SE-15 (Sigma). An autoclaved 1/50 concentration stock of the antifoam was added aseptically to the filter-sterilized (0.22µM) MHB/DMSO mixture. The only difference between the base medium and drug-containing medium was the addition of filter-sterilized Ciprofloxacin (Sigma-Aldrich). Starting populations of *E. coli* BW25113, *A. baumannii* ATCC 17978, and *P. aeruginosa* ATCC 27853 strains began from original glycerol stocks of parent organisms from ATCC; stocks were streaked onto LB-agar plates for isolation and grown overnight in MHB at 37°C. Liquid cultures were inoculated from single colonies and grown at 37°C, shaking to an optical density of 0.2-0.4 OD600, then diluted to 0.02 OD600 with drug-free medium in reactors to begin the run. All reactors in a single evolutionary run with *E. coli* were started from a liquid culture sourced from one isolated colony. Different individual colonies were chosen to seed reactor cultures for *A. baumannii* and *P. aeruginosa.* Glycerol stocks from these starting cultures were preserved, and cell pellets were used to determine genomic sequences to account for the anticipated larger diversity of preexisting variants (see below). Five separate evolutionary runs were performed: two with *E. coli* BW25113 (CEC-2 and CEC-4), one with *A. baumannii* ATCC17978 (CAB-1), and two with *P. aeruginosa* ATCC27853 (PAC-1 and PAC-2). Of the two *E. coli* runs, CEC-2 was performed under a steeper increase in drug concentration than CEC-4. This was achieved by using a higher drug concentration in Pump 2 media: 0.156 mg/L or 10-fold of CIP MIC value (0.0156 mg/L as determined for the unevolved strain in the same media) was used in the beginning of the run, increasing to 0.468 mg/L (30xMIC) on Day 2 of evolution and then to 2.34 mg/L (150xMIC) on Day 4. The second *E. coli* run (CEC-4) used a CIP gradient starting from the lower concentration of 0.001mg/L (0.625xMIC) and gradually increasing up to 1.248 mg/L (80xMIC) over a more extended time period (6 days). The CAB-1 run started at 0.195 mg/L (1.25xMIC, given MIC=0.156 mg/L for *A. baumannii* ATCC 17978 unevolved strain) and progressed up to 40x in 4 days. For morbidostat runs with *P. aeruginosa*, a modified control software was used (https://github.com/sleyn/morbidostat_v2_construction), which allows the user to vary the volume of drug-containing media added at every dilution step. Therefore, the media in Pump 2 remained at the concentration of 7.8mg/L (25-50xMIC, as for unevolved *P. aeruginosa* ATCC 27853 strain MIC=0.156-0.313 mg/L) throughout the entire experiment in both PAC-1 and PAC-2 runs, with a continual increase in drug concentration controlled through the software alone.

Over the course of all runs, dilutions were performed with V = 4 mL (20% of the reactor volume). The calculated changes in drug concentration in each reactor were plotted (as in **Figure 1c-e**) along with recorded changes in OD (see **Supplementary Methods** and **Supplementary Figure S2** for a complete set of parameters and plots generated for each run). As a result of optimization of the originally published morbidostat device and workflow (28), all runs were performed continuously (up to 6 days) without loss of sterility in bottles and feedlines. In the case of *P. aeruginosa* (but not *E. coli* or *A. baumannii*), the process included daily transfer of the culture (along with sample collection) to a fresh sterile glass reactor tube in order to minimize laser interference from biofilm gradually accumulating on the walls of the reactor. Otherwise, 10 mL samples were collected with a fresh sterile syringe via a dedicated needle port at one (or two) timepoints each day based on the OD, growth rates, and drug concentrations established in each reactor. All collected samples were used to prepare glycerol stocks (for further clonal analysis) and to extract genomic DNA from frozen cell pellets (prepared from the main portion of each sample).

### Genomic DNA extraction and sequencing

DNA was extracted using GenElute™ Bacterial Genomic DNA Kit Protocol NA2110 (Sigma Aldrich) according to the protocol for Gram-negative cells. Total DNA from evolutionary run samples were extracted from frozen cell pellets. DNA of selected clones was extracted from fresh pellets of liquid cultures grown 8–16 hrs at 37°C.

Nonamplified DNA libraries for Illumina sequencing of all population samples (and some of the analyzed clones) were prepared using NEBNext® Ultra™ II FS DNA Library Prep Kit for Illumina modules E7810L and E7595L (New England BioLabs) following the manufacturer protocol for use with inputs ≥ 100 ng with modifications to eliminate PCR-amplification steps.

IDT for Illumina TruSeq DNA UD Indexes 20022370 or TruSeq DNA CD Indexes 20015949 (Integrated DNA Technologies, Illumina) were used in place of NEBNext adaptors. Library size selection and clean up were performed using AMPure XP beads (Beckman Coulter) following the NEB protocol. An alternative faster and more cost-efficient approach was used for up to 96-plex sequencing of DNA from some of the analyzed clones. Clone DNA was prepared for Illumina sequencing using the PlexWell PW384 kit and included adaptors (seqWell) following manufacturer instructions.

Prepared libraries were quantified using NEBNext® Library Quant Kit for Illumina® E7630L (New England BioLabs) and pooled with volumes adjusted to normalize concentrations and provide for ∼1,000-fold genomic coverage for population samples (20–30 samples per HiSeq lane depending on the genome size) or ∼200-fold coverage (up to 96 samples per lane) for clones. Library size and quality were analyzed with the 2100 Bioanalyzer Instrument (Agilent). Pooled DNA libraries were sequenced by Novogene Co. on Illumina HiSeq X Ten or HiSeq 4000 machines using paired end 150bp read length.

Nanopore long-read sequencing was used to verify and complete assemblies for unevolved genomes. DNA samples were prepared for long-read Nanopore sequencing using the Nanopore Rapid Barcoding gDNA Sequencing kit SQK-RBK004 (Oxford Nanopore Technologies) according to the manufacturer protocol and sequenced using MinION and flow cell FLO-MIN106.

### Sequence data analysis and variant calling

The depiction of the sequencing data analysis pipeline (**Supplementary Figure S3**) and statistics for each sample (coverage, number of reads, and percentage of mapped reads) are provided in the Supplementary Materials (**Supplementary Table S1)**. Briefly, Illumina sequencing read quality was assessed with FastQC v. 0.11.8 (29). Adaptor trimming was performed using Trimmomatic v. 0.36 (30). In population sequencing data analysis, we used BWA MEM v. 0.7.13 for read alignment against reference genomes (31). SAM and BAM file manipulations were performed with PICARD v. 2.2.1 tools (https://broadinstitute.github.io/picard/) and samtools v. 1.3 (31). Realignment and base quality score recalibration were performed with the LoFreq Vitebri module (32) and Genome Analysis ToolKit v. 3.5 (33). For variant calling, SNPs and small indels were identified with LoFreq v. 2.1.3.1 (32). IS element rearrangements in population samples were predicted by a new iJump software developed for this purpose (https://github.com/sleyn/ijump). IS elements in reference genomes were predicted using the ISFinder database (34). Variant effects were annotated with snpEff v. 4.3 (35). VCF file manipulations were performed with bcftools v. 1.3 (36). Clonal sequencing data were analyzed using breseq software (37). Copy number variations were checked using the CNOGpro v. 1.1 R language package (38). R version 3.6.0 was used. Repeat regions were masked based on analysis produced by MUMmer v. 3.1 (39).

A *de novo* assembly of the *Acinetobacter baumannii* ATCC 17978 reference genomes for each of the six clones (samples A1–A6) used as the starting point for experimental evolution was accomplished by a hybrid approach combining Illumina (short reads) and Nanopore (long reads) data. Nanopore reads were base-called and demultiplexed using Albacore v. 2.3.4 base-caller (available on the ONT community site). To increase coverage with long reads, we performed a second round of demultiplexing of unclassified reads with Porechop v. 0.2.4 (https://github.com/rrwick/Porechop). Adaptors were trimmed with Porechop. Both reads called with Albacore and Porechop were combined and used along with Illumina reads to make *de novo* assembly with SPAdes v. 3.13.0 (40). The assembly was annotated using RASTtk web server (41). For detailed description of sequencing data processing see Supplementary Methods.

### Minimum inhibitory concentration (MIC) determination

MIC values were determined for parent strains and selected individual clones to connect mutations and genome rearrangements with resistance phenotype. MIC assays were prepared with twofold increasing concentrations of Ciprofloxacin by the broth dilution method following CLSI and EUCAST standard protocols, using resuspended fresh colonies in Cation-Adjusted Mueller Hinton broth medium (42). Measurements were performed using: (i) a growth curve method in microtiter plates at a wavelength 600nm in a BioTek ELx808 plate reader at 37°C (for *E. coli* and *A. baumannii*), or (ii) an end point analysis (for *P. aeruginosa,* results read at 17 hours).

### Data availability

Clonal and population sequencing data are available in the SRA database by BioProject accession number PRJNA598012 (https://www.ncbi.nlm.nih.gov/bioproject/PRJNA598012); the novel *A. baumannii* ATCC 17978 annotated assembly is available in the European Nucleotide Archive by sample accession number ERS4228590 (http://www.ebi.ac.uk/ena/data/view/ERS4228590). The reference genomes were downloaded from the PATRIC database(43). PATRIC IDs are: 679895.18 for *E.coli* BW25113, 287.6323 for *P. aeruginosa* ATCC 27853.

### Code availability

Custom code and detailed description of sequencing data processing is deposited to GitHub (https://github.com/sleyn/paper_cipro_EC_AB_PA_2021).

## Results and Discussion

### Evolution of CIP-resistance in *E. coli* BW25113

Two evolutionary runs were performed to assess the impact of different ranges and rates of CIP concentration escalation on the dynamics and spectra of acquired mutations in *E. coli* (**Supplementary Figure S2A, B**). For both runs, a complete list of significant sequence variants (passing all filters implemented in the computational pipeline) observed in each reactor and timepoint is provided in **Supplementary Tables S2B and S2C**.

Several preexisting low-frequency variants (mostly ∼2-3%) detected in the unevolved population at timepoint 0 were distinct between independently prepared inoculates used in each of the two runs, pointing to their stochastic nature. Most of these variants disappeared from populations over the course of selective outgrowth except two SNPs in genes encoding: (i) uncharacterized protein YigI:A146T (BW25113_3820); and (ii) uncharacterized transporter of BCCT family YeaV:V428D (BW25113_1801). These two variants expanded from 3% to 83% of the population in Reactor 2, and from 13% to 54% in Reactor 6, respectively, by the end of the CEC-2 run; notably, each mutation apparently coupled with the common GyrA:D87Y mutation (**Supplementary Table S2A)**.

The range and dynamics of major acquired mutations were generally similar in both runs, CEC-2 and CEC-4 (**Supplementary Tables S2A, S2B**). Of those, the earliest and the most prominent were missense mutations in the A and B subunits of DNA gyrase (GyrA and GyrB), a primary CIP target. These mutations typically emerged within the first 24 hrs and sustained or further expanded in all populations unless outcompeted by other primary target mutations. As such, a GyrB mutation Ser464Phe, which emerged at an early stage in all six reactors of the CEC-2 run, was rapidly outcompeted by GyrA mutant variants in three of these reactors (**Figure 2a, Supplementary Figure S4A**). An alternative GyrB mutant variant, Ser464Tyr, dominated all four reactors at an early stage of the CEC-4 run. This variant sustained in all but one reactor (**Supplementary Figure S4B**), where it was outcompeted by a GyrA:Asp87Tyr variant combined with a disruptive deletion in SoxR (**Figure 2b**). These distinctive evolutionary trajectories may be driven by a number of factors including different drug escalation profiles (**Supplementary Figure S2A,B**) and different effects of the two alternative substitutions at GyrB:Ser464 on CIP resistance and fitness.

**Figure 2.**
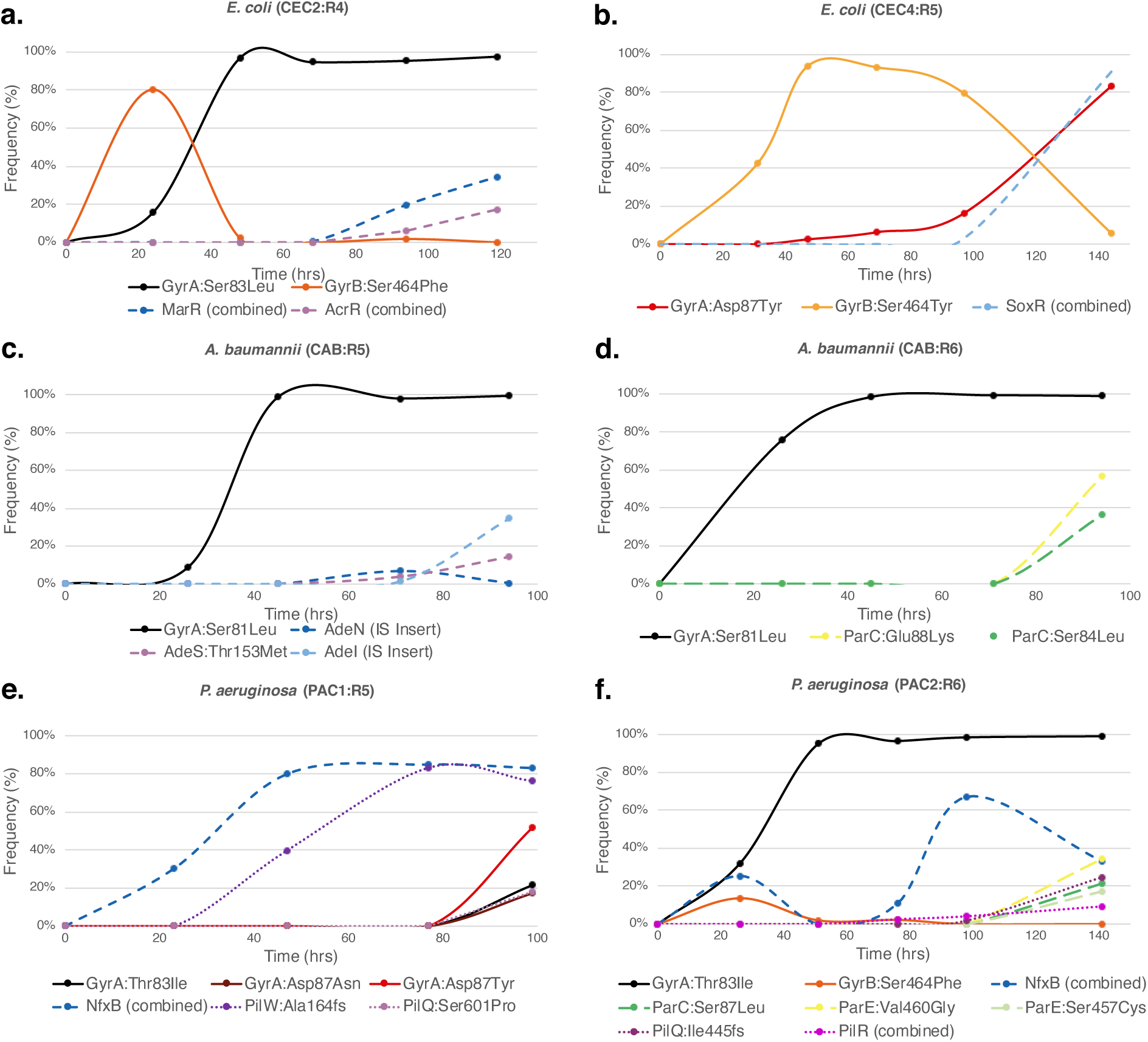
**Population dynamics of experimental evolution of ciprofloxacin resistance in *E. coli* (a, b)*, A. baumannii* (c, d), and *P. aeruginosa* (e, f).** Frequency of major mutations (reaching ≥5%) in evolving bacterial populations is shown as function of time for selected reactors. Selected reactors are shown: a) reactor 4 from CEC-2; b) reactor 5 from CEC-4; c) reactor 5 from CAB; d) reactor 6 from CAB; e) reactor 5 from PAC-1; f) reactor 6 from PAC-2.

The mutant variant GyrA:Asp87Tyr was among the most prominent variant in GyrA and the only common one between the two *E. coli* evolutionary runs. Other substitutions at this position included one major variant, Asp87Gly (observed at varying frequencies in all but one reactor of the CEC-2 run) and one minor variant, Asp87Asn (observed at low frequency in one reactor of the CEC-4 run). The only other high frequency GyrA:Ser83Leu variant was observed in one reactor (R4) of the CEC-2 run. These two residues are known to be the most commonly mutated *gyrA* residues in CIP-resistant *E. coli* (10, 44). Of the two other low frequency GyrA variants, the Ala119Glu substitution was previously reported in quinolone-resistant *E. coli* and *Salmonella* (10, 45). The second variant, GyrA:Asp82Gly, was also previously observed in these species as well as in CIP-resistant *Bartonella bacilliformis* (45, 46). All observed amino acid substitutions were located in the vicinity of the known CIP binding site, close to a DNA binding pocket at the interface of the DNA gyrase subunits A and B (**Figure 3**). No mutations in the Topoisomerase IV subunit A gene *parC,* a known secondary target of CIP, was observed in our study of *E. coli*, and only one low frequency variant with Arg303Ser mutation was observed in the gene *parE* encoding subunit B of this enzyme.

**Figure 3.**
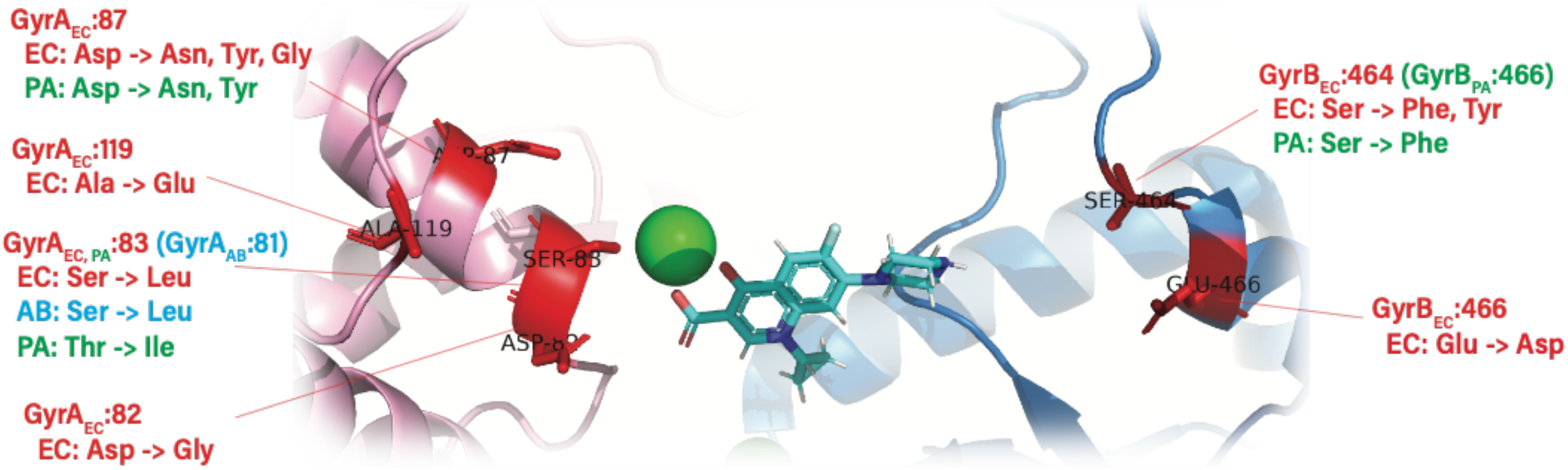
**Amino acid substitutions in GyrA (pink chain)/GyrB (blue chain) observed in the course of morbidostat-based experimental evolution of CIP resistance in *E. coli* (red font), *A. baumannii* (blue font), and *P. aeruginosa* (green font) mapped on 3D structure [PDB: 6RKW]**. The ciprofloxacin molecule (blue) and Mg^2+^ ion (green) were added by structural alignment of 6RKW with the structure of Mycobacterium tuberculosis gyrase bound to CIP (PDB:5BTC). The substitution equivalent to *P. aeruginosa* GyrB:Leu128Pro is not shown. It is located in proximity to the ATP-binding site of GyrB. Chain A of 6RKW was aligned to chain A of 5BTC using FATCAT. The same rotation-translation was then applied to all chains in 6RKW to align the full structure.

Clones representing major GyrA:Ser83Leu and GyrB:Ser464Phe/Tyr mutant variants without any additional mutations were isolated from respective reactors and exhibited an 8–16-fold increase in CIP-MIC values compared to the parental (unevolved) strain (**Supplementary Table S3A**). This magnitude of the effect on CIP-resistance is consistent with previous reports(44, 47). Nearly all isolated clones representing the GyrA:Asp87Gly variant contained additional mutations leading to efflux upregulation (as described below). In our study, GyrB mutations appear to have a somewhat smaller impact on resistance compared to GyrA mutations (on average ∼2-fold) as seen previously(48).

Numerous additional mutations emerged on the background of GyrA or GyrB mutant variants at a later stage of experimental evolution, dominated by frameshifts, disruptive deletions, and IS element insertions. These mutations appeared along with an increase of drug concentration, and most of these events are predicted to lead to upregulation of efflux machinery. The most prominent in both runs were the loss-of-function events affecting transcriptional regulators MarR, AcrR, and SoxR that negatively control the expression of well-known efflux pumps MarAB and AcrAB, and an outer membrane protein TolC (**Figure 4a**). Disruptive mutations occurred in over 30 distinct locations in the gene *marR* (**Supplementary Tables S2A,B**). Similar events were shown previously to increase MarA expression, thus contributing to CIP-resistance and to a broader multi-drug resistance phenotype(49, 50) . Clinical *E. coli* isolates with fluoroquinolone resistance commonly contain mutations in *marR*, causing constitutive expression of the *mar* operon (51). Deletion of the C-terminus of MarR increased *E. coli CIP* resistance *in vivo* (52), and inactivation of MarR has been shown to increase *marA* expression, effecting drug efflux by way of transcriptional amplification of *acrAB* and *tolC* pumps (17, 53).

**Figure 4.**
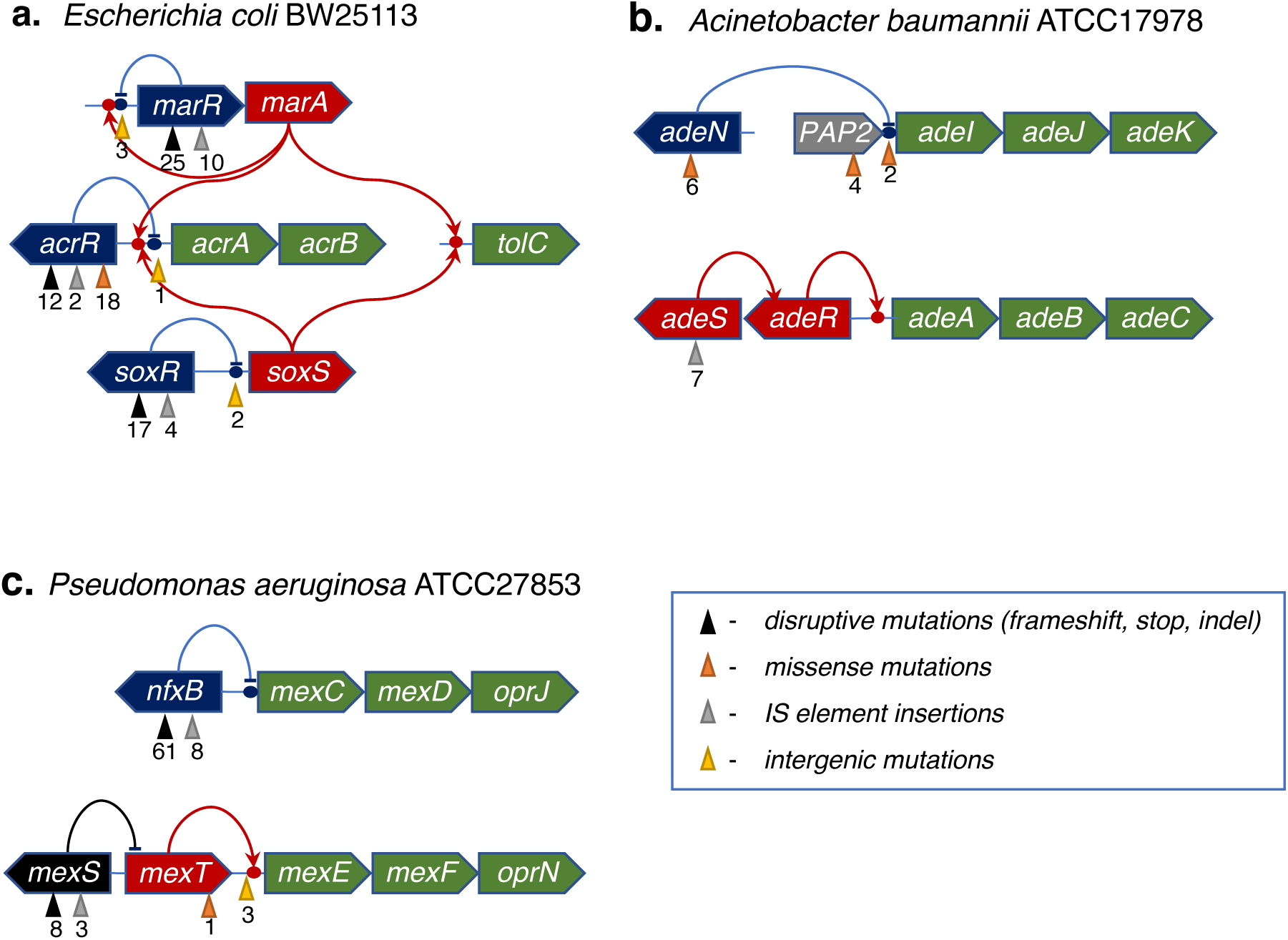
**Mutations leading to upregulation of efflux machinery in *Escherichia coli* (a), *Acinetobacter baumannii* (b), and *Pseudomonas aeruginosa* (c) detected over the course of experimental evolution of CIP resistance.** The total number of distinct variants detected in at least one of the reactors is shown under the color-coded upward triangles indicating a type of mutation

Among several missense mutations (observed only in CEC-2 run), the most prominent amino acid substitution MarR:Thr39Ile (which expanded up to 1/3 of bacterial population when combined with GyrA:Asp87Tyr in reactor R2) affected the known DNA-binding site of the MarR repressor (54). Dynamics of acquisition of MarR mutational variants appear to have been faster in the CEC-2 run than in CEC-4 (after 48 hrs vs 96 hrs, respectively) reflecting the sharper increase of drug concentration in CEC-2 run. Clones with *marR* mutations combined with *gyrA* or *gyrB* mutations exhibited MIC of 4–8-fold higher than corresponding clones with only mutations in DNA gyrase (**Supplementary Table S3A**).

IS element insertions, frameshifts, and deletions occurred in multiple positions of both the *acrR* gene encoding a transcriptional regulator of the *acrAB* operon, as well as in the intergenic region of the *acrA<>acrR* divergon (**Figure 4a**). These variants appeared at relatively low frequency at the latest stage of experimental evolution, after ∼90 hrs in both runs (**Figure 2a, Supplementary Figure S4A,B**). Among isolated clones, *acrR* mutations were found only as combined with mutations in *marR* and one of the DNA gyrase subunits (*gyrA* or *gyrB*). These triple mutants exhibited the highest MIC values observed in this study, up to 128-fold higher than wildtype MIC (**Supplementary Table S3A**).

Point mutations, frameshifts, and deletions also occurred frequently in *soxR*, the redox-sensitive repressor which negatively regulates SoxS, transcriptional activator of efflux genes *acrAB* and *tolC* (**Figure 4b**). The SoxR variant Gly121Asp, which occurred in the CEC-2/R5 population, has recently been shown to increase SoxS expression 12-fold (50). Similar to *marR*, mutations in *soxR* emerged between 48 and 96 hrs, more rapidly in CEC-2 than CEC-4 (**Figure 4b, Supplementary Figure S4A** and **Supplementary Figure S4B**). Deletions and small insertions in *soxR* were found in clones along with DNA gyrase mutations contributing to a further increase of MIC by a factor of 4–8x (**Supplementary Table S3A**).

Despite the relatively low abundance of individual mutations in efflux regulators *marR*, *acrR*, and *soxR*, a very high combined frequency of efflux-upregulating mutations in both CEC-2 and CEC-4 makes increased efflux a predominant mechanism for elevating CIP-resistance on the background of *gyrAB* mutations. This trend is consistent with other published studies including laboratory evolution experiments and analysis of CIP-resistant clinical isolates of *E. coli* (23, 55).

### Evolution of CIP-resistance in *Acinetobacter baumannii* ATCC 17978

To accurately assess preexisting variations that occur at much higher frequency in *A. baumannii* than in *E. coli* K12, we first sequenced and assembled a complete genome corresponding to our stock of *A. baumannii* ATCC 17978 (ENA Project: PRJEB36129). This assembly was further used as a reference for this study and featured substantial differences with publicly available sequences: 16 variants compared to the GCA_001593425 assembly and 87 variants compared to the GCA_000015425.1 assembly (at >85% frequency threshold) over 98.5% and 99.0% mapped reads, respectively. Unlike in *E. coli* runs, we prepared each of the starter cultures from individual colonies and sequenced total genomic DNA isolated from these cultures (samples A1-A6). To account for potential genomic rearrangements, we used a hybrid approach combining the data from high-coverage Illumina sequencing (short reads) with Oxford Nanopore sequencing (long reads). In contrast to the public genomes, this approach revealed that all starting cultures have pAB3 (56) as an extrachromosomal plasmid (in public genomes it is integrated to the chromosome) and an additional 52kb locus (**Supplementary Table S1C**).

The single *A. baumannii* evolutionary run was performed using a mild drug escalation regimen which started from a CIP concentration in Pump 2 corresponding to 1.25xMIC. The dynamics of CIP-resistance acquisition observed in five parallel reactors appeared faster in the case of *A. baumannii* as compared to *E. coli* (**Figure 1d, Supplementary Figure S2C**). One reactor (R1) was excluded due to technical failure. Resistance was driven at the first stage by a single *gyrA* mutation Ser81Leu, which was first detected after Day 1 and expanded to ≥95% by Day 3 (**Figure 2c,d, Supplementary Table S2D**). This mutation leads to ∼8-fold increase in MIC (**Supplementary Table S2B**), which is similar to the impact of the position-equivalent GyrA:Ser83Leu variant of *E. coli* (**Supplementary Table S2A**). The same mutation is commonly found in CIP-resistant *A. baumannii* isolates with a similar impact on MIC, as reported for *A. baumannii* ATCC 19606 (57).

At the next stage of experimental evolution, adaptation to higher drug levels was attained via a variety of mutations apparently upregulating drug efflux machinery (**Figure 2c, Figure 4b, Supplementary Figure S4C, Supplementary Table S2C**). Such upregulation was achieved with three classes of variants: (i) inactivation of transcriptional repressor AdeN of the efflux pump AdeIJK caused exclusively by insertion of different mobile elements (IS*Ab1*, IS*Ab11*, and IS*Ab12*); (ii) polar effects from insertion of IS elements (IS*Ab1*, IS*Ab12*, IS*Ab18*, and IS*Abcsp3*) in the intergenic region or in the gene encoding membrane-associated phospholipid phosphatase immediately upstream of the *adeI* gene; and (iii) several missense mutations in the AdeS sensory histidine kinase of a two-component system (AdeSR) positively regulating another efflux pump AdeABC (58–60). IS element insertions upstream of the *adeIJK* operon were previously reported to reduce susceptibility of *A. baumannii* to ciprofloxacin and other substrates of the respective efflux pump (61, 62). Notably, one of the AdeS mutant variants, AdeS:Asp60Gly, expanded up to 65% in reactor R3 population by the end of Day 4. An apparent prevalence by abundance of AdeABC vs AdeIJK upregulating variants (see Table 1) is consistent with a recent report on higher efficiency of the AdeABC transporter than the AdeIJK transporter in efflux of hydrophilic agents such as fluoroquinolones (63).

**Table 1.** Major mutational variants detected in the course of experimental evolution of ciprofloxacin resistance^a^

| *E. coli* |  |  | *A. baumannii* |  |  | *P.aeruginosa* |  |  |
| --- | --- | --- | --- | --- | --- | --- | --- | --- |
| Variant | Max. Freq. (%)<sup>b</sup> | In reactors (out of 10) | Variant | Max. Freq. (%)<sup>b</sup> | In reactors (out of 5) | Variant | Max. Freq. (%)<sup>b</sup> | In reactors (out of 12) |
| **Primary targets** |  |  |  |  |  |  |  |  |
| **GyrA, DNA gyrase subunit A** |  |  |  |  |  |  |  |  |
| Ser83Leu | 98% | 1 | Ser81Leu | 99% | 5 | Thr83Ile | 100% | 12 |
| Asp87Asn | 17% | 1 |  |  |  | Asp87Asn | 79% | 2 |
| Asp87Tyr | 99% | 4 |  |  |  | Asp87Tyr | 52% | 1 |
| Asp87Gly | 74% | 6 |  |  |  |  |  |  |
| Ala119Glu | 7% | 2 |  |  |  |  |  |  |
| **GyrB, DNA gyrase subunit B** |  |  |  |  |  |  |  |  |
| Ser464Phe | 96% | 6 |  |  |  | Leu128Pro | 7% | 1 |
| Ser464Tyr | 99% | 4 |  |  |  | Ser466Phe | 63% | 4 |
| **Secondary targets** |  |  |  |  |  |  |  |  |
| **ParC, Topoisomerase IV subunit A** |  |  |  |  |  |  |  |  |
|  |  |  | Ser84Leu | 36% | 4 | Ser87Leu | 53% | 6 |
|  |  |  | Glu88Lys | 86% | 2 |  |  |  |
|  |  |  | Asn334Tyr | 4% | 1 |  |  |  |
| **ParE, Topoisomerase IV subunit B** |  |  |  |  |  |  |  |  |
|  |  |  |  |  |  | Leu121Pro | 8% | 1 |
|  |  |  |  |  |  | Ser457Gly | 6% | 1 |
|  |  |  |  |  |  | Ser457Cys | 17% | 1 |
|  |  |  |  |  |  | Ser457_insArg | 9% | 1 |
|  |  |  |  |  |  | Val460Gly | 62% | 2 |
| **Efflux pump deregulation** |  |  |  |  |  |  |  |  |
| *marR*<sup>c</sup> | 86% | 9 | *adeN*<sup>c</sup> | 28% | 5 | *nfxB*<sup>c</sup> | 90% | 12 |
| *soxR*<sup>c</sup> | 95% | 9 | *adeS*<sup>c</sup> | 72% | 4 | *us_mexE* | 39% | 1 |
| *us_soxS* | 10% | 2 | *us_adel* | 36% | 4 | *us_mexT* | 12% | 5 |
| *acrR*<sup>c</sup> | 63% | 9 |  |  |  | *mexT:G58D* | 6% | 1 |
| *us_acrA* | 53% | 3 |  |  |  | *mexS*<sup>c</sup> | 94% | 7 |
| **Other strongly implicated genes** |  |  |  |  |  |  |  |  |
|  |  |  |  |  |  | *pilBOPQRSTWZ*<sup>c</sup> | 98% | 11 |
|  |  |  |  |  |  | *mutL*<sup>c</sup> | 37% | 2 |
|  |  |  |  |  |  | *mutS*<sup>c</sup> | 27% | 3 |

<sup>a</sup>Listed mutations that reached $\geq 5\%$ in at least in one reactor and one time point.
<sup>b</sup>Maximum observed frequency reached at any time point in any reactor.
<sup>c</sup>Disruptive mutations (mainly frameshifts, stops, small indels and IS inserts).

The upregulation of the efflux pump AdeIJK on the background of the GyrA:Ser81Leu variant led to an additional 2–4-fold increase in MIC (**Supplementary Table S3B**). This was determined for individual clones representing mutant variants of both types: IS-insertions in the *adeN* gene, and IS-insertions in the intergenic region upstream of gene *adeI* (16xMIC and 32xMIC, respectively compared to unevolved strain). No mutations were observed in genes associated with yet another known multidrug efflux system of *A. baumannii*, AdeFGH (64).

Finally, the highest level of CIP-resistance was achieved as a result of a combination of the initial GyrA:Ser81Leu variant with additional missense mutations in the *parC* gene encoding DNA topoisomerase IV subunit A, a secondary target of fluoroquinolone drugs. These mutations emerged only at the latest stage of experimental evolution (Day 4)(**Figure 2d, Supplementary Figure S4C**). The variant ParC:Asn334Tyr only reached low frequency (4% in R2) in one reactor, whereas two other variants (ParC:Ser84Leu and ParC:Glu88Lys) emerged in several reactors reaching much higher frequency (up to 86%)(**Supplementary Table S2C**). Both variants exhibited resistance of 128-fold compared to the unevolved strain, the highest increase in MIC observed with individual clones (**Supplementary Table S2B)**. These two mutated residues are position-equivalent to the most commonly mutated Ser80 and Glu84 residues in *E. coli* ParC (57, 65, 66). A commonly reported CIP-resistant mutation Ser458Ala in the Topoisomerase IV subunit B gene *parE* (67) was observed in the last timepoint in only one reactor (R1) at low abundance (4%).

### Evolution of CIP-resistance in *Pseudomonas aeruginosa* ATCC27853

To further expand the comparative resistomics approach and assess common and species-specific trends in the dynamics of acquisition and mechanisms of CIP-resistance, we applied the optimized morbidostat-based experimental evolution workflow to study another important Gram-negative pathogen, *Pseudomonas aeruginosa*. Two evolutionary runs (PAC-1 and PAC-2) with *P. aeruginosa* ATCC27853 were performed using starter cultures prepared from six distinct colonies. In contrast to *A. baumannii*, this analysis did not detect any substantial variations between these cultures (**Supplementary Table S2D, S2E**). Some of the starter cultures contained up to 20 preexisting low frequency variants (in a range of 1-10%). These low-frequency variants reflect stochastic microheterogeneity, and they typically disappeared from bacterial populations after Day 1 of selective outgrowth in the morbidostat (see **Supplementary Table S2D, S2E**).

A technical challenge originated from a tendency of *P. aeruginosa* to form a visible biofilm on the glass surface of the reactor, especially located near the interface with air. This problem was mitigated by keeping culture densities well within logarithmic growth phase (OD600≤0.55) and by daily transfer of cultures to clean reactors, which was performed along with sample collection.

Using an improved version of morbidostat software allowed us to perform two evolutionary runs with more flexible iterative modulation of drug concentration during the runs. The second run (PAC-2) employed a shallower drug escalation mode in the early stage (Days 1-3) and enhanced the overall duration of the experiment (6 days instead of 4 days in PAC-1) with a larger number of collected and analyzed samples (**Supplementary Figure S4D,E).** In contrast to the case of *E. coli*, varying critical parameters of morbidostat runs with *P. aeruginosa* appeared to have some notable impacts on the range and dynamics of acquisition of certain mutational variants as outlined below (**Figure 2e,f, Supplementary Figure S4D,E**).

The most prominent primary mutation appearing in all reactors of both the PAC-1 and PAC-2 runs was the GyrA:Thr83Ile variant (**Table 1**), which is equivalent to GyrA:Ser83Leu in *E. coli* and Ser81Leu in *A. baumannii*. This was the only GyrA variant in the PAC-2 run, and it emerged typically on Day 2 (in four out of six reactors) and expanded up to ∼100% of population by Day 3-4 in all reactors. The same GyrA:Thr83Ile variant was also universally present and dominant in all six reactors of the PAC-1 run. However, it appeared at a somewhat later stage (typically on Day 3), in most cases after disruptive mutations in the *nfxB* gene (see below). Additional variants, GyrA:Asp87Asn and GyrA:Asp87Tyr, appeared in two reactors of the PAC-1 run and partially outcompeted the GyrA:Thr83Ile variant by the end of the run (**Figure 2e,f, Supplementary Figure S4D,E**). Two of these three variants (GyrA:Thr83Ile and GyrA:Asp87Asn) represent the two most commonly mutated positions in CIP-resistant *P. aeruginosa* reported for both clinical and laboratory isolates (68, 69).

The only prominent GyrB:Ser466Phe variant (equivalent of the GyrB:Ser464Phe variant observed in *E. coli*) was detected transiently in one reactor in each run (R6 of PAC-1 and R5 of PAC-2), peaking at 30% and 63% of respective populations only to be entirely outcompeted by GyrA:Thr83Ile-containing variants by the end of both runs.

Among the most striking differences observed between the two runs was the appearance of mutations in the genes *parC* and *parE* encoding both subunits A and B of DNA topoisomerase IV exclusively in PAC-2. Remarkably, the most prominent ParC:Ser87Leu variant appeared in all six reactors on the last day of the run when the drug concentration was highest, reaching up to 53% in population as a secondary mutation on the background of GyrA:Thr83Ile-containing variants (**Figure 2f, Supplementary Figure S4E**). Among more diverse (albeit less universal) ParE variants, the most prominent were ParE:Val460Gly and ParE:Ser457Cys, both reaching the highest abundance (34% and 17%, respectively) in the same reactor (R6) on the last day of the PAC-2 run (**Figure 2f**). These exhibited the same general evolutionary dynamics as ParC variants, and also emerged on the background of the GyrA:Thr83Ile variant (**Figure 2f**).

Among the mutational variants driving efflux upregulation, the most common and abundant were various types of disruptive mutations in the *nfxB* gene encoding a transcriptional repressor of the *mexCD–oprJ* operon (70, 71)(**Figure 4e,f**). Disruptive mutations in the *nfxB* gene were reported to have pleiotropic effects improving fitness and antibiotic resistance, contributing to lowered expression of outer membrane porins and improved drug efflux (72, 73). In our study, the entire range of 61 mutations unambiguously leading to a loss of function (frameshifts, nonsense mutations, and indels) and 8 missense mutations were found spread over all 12 reactors in both PAC runs collectively reaching from 35% to 90% abundance in at least one time point in every reactor (**Supplementary Figures S4D,E**). However, the dynamics of their appearance and accumulation was strikingly different between the two runs. Indeed, in nearly all reactors of the PAC-2 run (except R3), these mutations emerged and accumulated after or together with the driving GyrA:Thr83Ile variant, similar to the evolutionary dynamics patterns observed for *E. coli* and *A. baumannii*. In contrast, in all reactors of the PAC-1 run, NfxB mutational variants reached > 30% overall abundance prior to comparable accumulations of GyrA variants that emerged later and expanded on the background of NfxB and/or other adaptive mutations (**Supplementary Figure S4D, Supplementary Table S2D).** This observation provides another example of how the difference in drug escalation regimen may affect evolutionary trajectories of CIP-resistance in *P. aeruginosa*.

Most of the individual NfxB disruptive variants in the PAC-2 run were present at relatively low frequency (2 – 20%) and randomly distributed among reactors. Remarkably, one NfxB variant, a substitution of the stop codon by a cysteine codon, which resulted in extension of the NfxB protein by 68 amino acids, appeared in all six reactors on the background of the GyrA:Thr83Ile variant, peaking at up to 85% abundance in population (**Supplementary Table S2E**). In every case, the abundance of this variant shrank to <20% on the last day of the run. This mutation was characterized earlier and found to lead to substantial overexpression of the MexCD-OprJ efflux pump (74). A similar, but less contrasted picture was observed in the course of the PAC-1 run in which the same variant appeared transiently in the middle of the run in five out of six reactors reaching 10-20% abundance, after which it completely disappeared from populations by the end of the run (**Supplementary Table S2D**). The single most abundant disruptive NfxB:Ile65fs variant reached >80% frequency in population and sustained until the end of the run, coupling with at least two out of three GyrA variants that emerged in this reactor (R5) only on the last day of PAC-1 run.

Additional low frequency mutations potentially upregulating another multidrug efflux pump MexEF-OprN (75) were detected in the upstream region of the respective operon (typically at a later stage, see **Supplementary Table S2D, S2E**). One of these was a missense mutation (Gly258Asp), and two were intergenic mutations in the upstream region of the gene *mexT* encoding its transcriptional activator (**Figure 4c**). However, the largest variety of mutations reaching up to 85% in all but one reactor in PAC-1 (and one reactor in PAC-2) were found within the coding sequence of a gene encoding an uncharacterized oxidoreductase MexS located in the divergon with MexT, immediately upstream of *mexE*, the first gene of the *mexEF-oprN* operon (**Figure 4c**). It was previously shown that a transposon inactivation of the *mexS* gene leads to upregulation of this typically quiescent operon via a yet unknown mechanism (76). Single amino acid substitutions in MexS leading to MexT-driven activation of the *mexEF-oprN* operon are frequently found in clinically isolated *nfxC* mutants of *P. aeruginosa* (a general term for strains with upregulated MexEF–OprN efflux pump) displaying enhanced virulence and drug resistance (77). Both MexCD–OprJ and MexEF–OprN systems affected in this study are known to be primary drivers of fluoroquinolone resistance; no mutations were found in several other known efflux systems of *P. aeruginosa* (78).

The last type of frequent mutations (including frameshifts, indels, and IS inserts) was observed at the late stage of both PAC-1 and PAC-2 evolutionary runs in several *pil* genes involved in Type IV pilus and fimbria biogenesis/assembly (**Table 1, Supplementary Table 2D,E**). Some of these mutations expanded to high abundance when coupled with GyrA mutational variants, e.g. up to 88% for PilQ:Gln232fs (R1 in PAC-1) and 65% for PilS:Gln14fs (R4 in PAC-2). Notably, a mutant variant PilW:Ala164fs appeared (in R5 of PAC-1) on Day 2 on the background of the NfxB:Leu62fs variant (in the absence of any GyrA/B mutations). The double mutant NfxB:Leu62fs/PilW:Ala164fs expanded to ∼85% of the population on Day 3 and provided a genetic background for the appearance of the three GyrA mutant variants on Day 4 (**Figure 2c, Supplementary Table 2D)**. The loss of type IV pili has been observed previously under CIP stress in *P. aeruginosa* (79). While the mechanistic rationale for this class of events remains unclear, it was hypothesized that the loss of type IV pili, known receptors for filamentous phages implicated in chronic infection, contributes to resistance against superinfection and lysis under ciprofloxacin stress (80).

The analysis of representative clones selected from PAC-1 confirmed the existence of CIP-resistant NfxB mutational variants lacking target-based mutations. Such clones exhibited about 8- fold increase in MIC. A combination of target-based (GyrA/GyrB) mutations combined with the loss of NfxB and/or other adaptive mutations discussed above increased the CIP-resistance up to 16–32-fold MIC as compared to the unevolved strain (**Supplementary Table S3C**). Given the observed complexity of *P. aeruginosa’s* pathways to CIP resistance, establishing the actual contribution and mechanistic effects of individual mutations would require additional studies.

Another distinctive feature observed in experimental evolution of *P. aeruginosa* was the emergence of disruptive mutations in *mutS* and *mutL* genes encoding DNA mismatch repair proteins. The appearance of MutS frameshift variants in R5 at the very end of PAC-1 run coincided with a spike of mutations in the same abundance range (**Supplementary Table 2D)**. More remarkable is a simultaneous occurrence of both mutational variants, MutS: Ala358fs (27%) and MutL:Leu706Arg (31%), in the same reactor (R4) on the last day of PAC-2 run. Despite similar abundances, these two variants likely represent two distinct clonal sub-populations, each accompanied by a broad range of secondary mutations (**Supplementary Table S2E**). Most of the accompanying secondary mutations did not occur at any other time points or in any other reactor. Loss of MutS function is known to increase the frequency of DNA replication errors leading to an explosion of mutations as demonstrated in many bacteria including *P. aeruginosa* (81). While limited to a single reactor per run, such a trajectory is not uncommon for *P. aeruginosa,* which was reported to acquire *mutS* loss-of-function mutations in cystic fibrosis patients possibly accelerating adaptation to the host environment and acquisition of antibiotic resistance (82). That said, the actual impact of *mutS*/*mutL* disruption on evolution of CIP resistance in our studies is not obvious. As already mentioned, MutS and MutL variants appeared only on the last day of each evolutionary experiment, whereas multiple co-appearing mutations seemed unrelated to drug resistance. Based on the abundance, only the GyrA:Asp87Asn variant (17%), which also emerged in R5 at the last time point of PAC-1 run, could have co-appeared on the background of the MutS:Arg302fs variant. However, even if confirmed by isolation of a corresponding double mutant (not accomplished in this work), this single event would not provide sufficient evidence of the hypothesized importance of a hypermutability phenotype to evolution of CIP resistance in *P. aeruginosa* which readily occurred in our study without any *mutS*/*mutL* disruption.

### *Comparative resistomics*: shared and unique features of evolutionary trajectories to CIP resistance in *E. coli*, *A. baumannii,* and *P. aeruginosa*

The present experimental evolution studies provided a foundation for comparative resistomics analysis of three representative Gram-negative bacterial species. Observation of each species in standardized continuous culturing conditions permitted discernment of common and species-specific aspects of evolutionary trajectories toward CIP resistance (**Figure 5**). The observed variety of these evolutionary trajectories can be approximated by a largely shared two-stage process. In Stage I, when the drug pressure was moderate (typically Day 1-3 of the morbidostat run), the emerging resistance (typically 4-16-fold MIC of unevolved parental strain) was usually driven by a single mutation rapidly expanding over the entire bacterial population in each reactor (**Figure 1c-e, Supplementary Figure S2**). In all reactors of *E. coli* (CEC-2 and CEC-4), *A. baumannii* (CAB-1), and one of the two morbidostat runs of *P. aeuruginosa* (PAC-2), the earliest (Stage I) mutations occurred in one of the two subunits of DNA gyrase (GyrA or GyrB). Among them, the most prominent and sustainable were amino acid substitutions in two positions of GyrA: Ser/Thr83 or Asp87 (by numeration of *E. coli* GyrA). At least one of these GyrA variants ultimately appeared in nearly all reactors expanding up to 100% abundance by the end of each run (**Table 1**, **Supplementary Table S2A-E**). Not surprisingly, these amino acid residues located in the CIP binding site (**Figure 3**) are the positions of the most common mutations in CIP-resistant clinical isolates (10, 44). Of these two positions, the former was the only one affected in *A. baumannii* (5 out of 5 reactors), and the most universal (in 12 out of 12 reactors) among the two affected positions in *P. aeruginosa* GyrA in our study. Amino acid substitutions at this position were shown to substantially reduce drug binding to the GyrAB:DNA complex (83) and thus confer a higher increase in MIC compared to other *gyrA* mutations (48). Notably, the GyrA:Ser83Leu variant appeared in only one out of 10 reactors of *E. coli* runs (**Table 1**), while most other reactors were dominated by GyrA:Asp87(Tyr, Gly, or Asn) variants.

**Figure 5.**
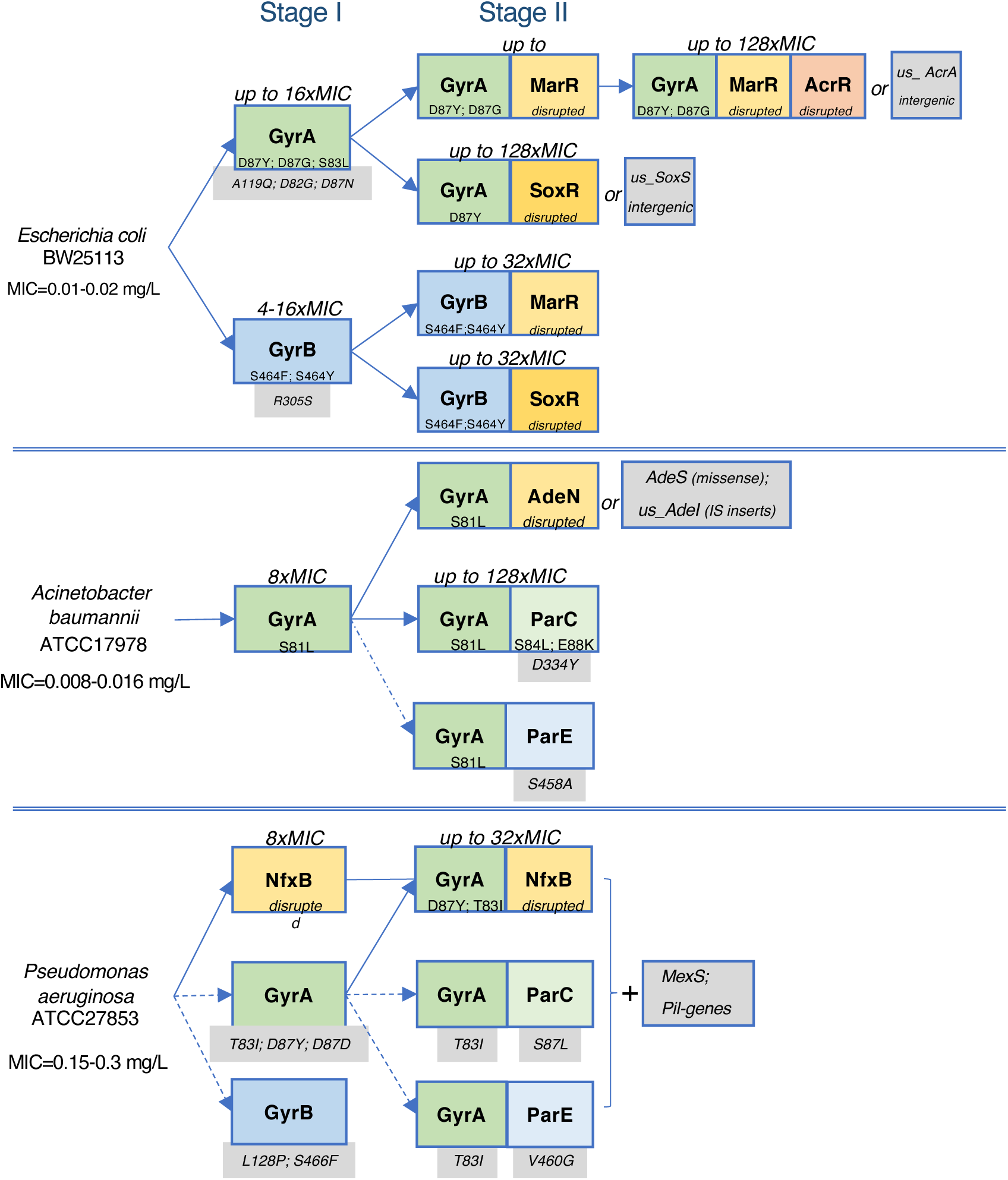
**Trajectories and stages in the experimental evolution of CIP resistance in *E. coli*, *A. baumannii*, and *P. aeruginosa.*** Major driver mutations are shown in color-coded boxes: missense mutations in targets (green and blue); disruptive mutations in efflux regulators (yellow and brown). Additional potentially relevant mutations detected in populations but not in isolated clones are shown on a grey background.

Another distinctive feature of evolution of CIP resistance in *E. coli* was a somewhat unexpected prominence of GyrB mutational variants. Thus, GyrB:Ser464Phe and GyrB:Ser464Tyr emerged as the earliest Stage I variants in all 6 reactors of CEC-2 and all 4 reactors of CEC-4 (**Table 1**, **Supplementary Figure S4AB**). Moreover, they sustained their predominant abundance until the end of both runs in 7 out of 10 reactors via coupling with efflux-deregulating mutations in Stage II. While GyrB is not considered a direct target of fluoroquinolones; the affected Ser464 position is located close to the CIP binding site in the GyrA:GyrB complex (**Figure 3**)(84). Further, CIP resistance-conferring mutations in this position were previously reported in *Citrobacter freundii*, *Morganella morganii*, *Salmonella typhimurium*, and *P. aeruginosa* (85–88). Notably, no GyrB variants were observed in *A. baumanni*, and a position-equivalent GyrB:Ser466Phe variant appeared only transiently in *P. aeruginosa* morbidostat runs (**Table 1**, **Supplementary Figure S4DE**).

Important similarities along with interesting differences can be deduced from the comparative analysis of mutational profiles of these three bacterial species with respect to a known secondary target of fluoroquinolones, DNA topoisomerase IV, comprised of two subunits (ParC and ParE). In contrast to *A. baumanniii* and *P. aeruginosa,* no mutations were detected in *parC* or *parE* genes in either one of the two *E. coli* morbidostat runs (**Table 1**). Moreover, even in the other two species, such mutations were relatively rare and occurred exclusively in Stage II on the background of Stage I-born GyrA mutations (**Supplementary Figure S4C-E)**. An interesting difference between these species is that while *A. baumanniii* featured only ParC variants, the second *P. aeruginosa* morbidostat run (PAC-2, but not PAC-1) revealed a substantial representation of both ParC and ParE variants (Table 1). Despite a clear distinction of mutational profiles and frequencies between the primary (GyrA/B) and secondary (ParC/E) CIP targets, the complete lack of ParC/E mutational variants in evolutionary experiments with *E. coli* is somewhat surprising given the reported presence of such mutations in ciprofloxacin-resistant clinical isolates of *E. coli* (67, 89). A rationale behind this single discrepancy between otherwise fully overlapping spectra of CIP target-directed mutations observed in morbidostat setting vs clinical isolates is unclear.

In addition to a narrow set of missense mutations in universal target genes, numerous different but mostly disruptive mutations emerged in a variety of efflux-regulating genes in all three species examined (**Table 1**, **Figure 4**). Among the common features of these mutational events: (i) they typically occurred in Stage II on the background of already accumulated GyrA/B variants, the universal evolutionary trajectory in *E. coli* and *A. baumannii* and predominant one in *P. aeruginosa*, and (ii) the most common targets of these mutations in all three species were negative regulators (transcriptional repressors) of efflux pump operons. Not surprisingly, a large fraction of such mutations includes nonsense mutations (stops), frameshifts, small indels, and IS insertions (**Figure 4**) representing a clear loss-of-function. Notably, the last of these was the most common, if not the only type of genetic alteration leading to loss of gene function in *A. baumannii.* In *E. coli*, IS insertions comprised more than half of the loss-of-function variants of negative efflux regulator AcrR (but not MarR or SoxR) and two intergenic variants potentially leading to derepression of the positive efflux regulator SoxS. Less frequent mutational events with the same type of downstream effects (upregulation of efflux pumps) occurred in intergenic regions (possible binding sites of respective transcriptional regulators). Additionally, several exclusively missense mutations arose in positive regulators (AdeS in *A. baumannii* and MexT in *P. aeruginosa*).

A notable deviation from the nearly universal evolutionary trajectory (target first, efflux later) was observed in *P. aeruginosa* for NfxB, a transcriptional repressor of the *mexCD-oprJ* efflux operon. NfxB-inactivating mutations emerged in both Stage I (in the absence of any target mutations; in PAC-1 but not in PAC-2) and Stage II of CIP resistance evolution in *P. aeruginosa*.

All major types of CIP resistance-conferring mutations described in the literature were observed in at least one of the three species in our study. These results support the utility of the established morbidostat-based workflow to elucidate antibiotic resistance mechanisms in a comprehensive manner. Strengthened further by a comparative resistomics approach, this study allowed us to elucidate and generalize major pathways to CIP-resistance in a group of divergent Gram-negative bacterial pathogens.

### Concluding remarks

We employed a comparative dynamic analysis of genetic adaptation to reveal both shared and distinctive features of three divergent Gram-negative bacterial species’ evolutionary trajectories towards CIP resistance. Despite obvious differences between the experimental conditions in the morbidostat and the complexity of processes leading to drug resistance in bacterial infections, the results obtained in this study are generally consistent with those deduced from clinical CIP-resistant isolates.

Many studies have suggested that resistance mutations observed in clinical specimens are biased toward low fitness costs, as fitness may be more important than the extent of resistance in clinical conditions (10, 55, 90). This observation is perhaps best exemplified by the apparent bias against inactivation in *marR* mutations in ciprofloxacin-resistant isolates. The interplay of fitness and resistance may explain the general mutation dynamics observed in this study. An initial emergence of rare target mutations with a presumably low fitness-cost occurred in the morbidostat similar to what is observed in the clinic. This was followed by an explosion of various efflux-regulator mutations later in the evolutionary trajectory as high drug resistance became the primary hurdle, and additional adaptation mechanisms had to be engaged for survival (10). Especially in human infection, early selection appears to favor the mutation causing the lowest fitness detriment, even when the resulting increase in resistance is relatively modest (55).

Indeed, disruptive mutations observed abundantly in efflux regulator genes in our study are statistically much more likely to occur than a few beneficial amino acid substitutions in a very limited set of positions in CIP target enzymes. And yet, such mutations were mostly observed as secondary events at a later stage (Stage II) of experimental evolution in the morbidostat. This is consistent with the observations that these mutations in clinical CIP resistant isolates also appear only in addition to target-directed mutations, likely due to relatively higher fitness costs. This overall consistency with data from clinical isolates is possibly driven by a relatively large bacterial population sizes maintained in the morbidostat setup, along with continuous competition at the level of growth rate (fitness) imposed by frequent dilutions. In contrast, more traditional experimental approaches are constrained by population-limiting bottlenecks which cause minimal competition and contribute to the selection of low fitness variants (23).

*Overall*, this study (as well as other similar studies (26–28, 91) confirms that morbidostat-based experimental evolution provides a powerful approach to assess the dynamics and mechanisms of antimicrobial resistance acquisition in a broad range of pathogens. This methodology is scalable and applicable for known antibiotics as well as novel drug candidates. The utility of comparative resistomics to assess and triage drug candidates across a range of target pathogens is expected to manifest even in the early phase of antimicrobial drug development. Combined with standard efficacy and safety evaluation, such assessment would contribute to the rational selection of compounds capable of providing lasting therapies in the field for longer periods of use.

## Acknowledgements

This work was supported by F. Hoffmann-La Roche Ltd pRED postdoctoral fellowship to S.L. and by Laboratory Funding Initiative of SBP to A.O.

## For reviewers

1. The sequencing data for population and clonal sequencing is available in NCBI SRA database through the reviewers login (https://dataview.ncbi.nlm.nih.gov/object/PRJNA598012?reviewer=ssd6539aliv2i9m8qs8g6qsc5k)
2. The manuscript “Experimental evolution in morbidostat reveals converging genomic trajectories on the path to triclosan resistance” by S. Leyn *et al*. that was accepted but not yet published is available by the following link: https://16515-my.sharepoint.com/:u:/g/personal/sleyn_sbpdiscovery_org/ERMAw0tAs8xPpOoRrhoGlF4Ba_bObjanBCr4oUSyAuxvag?e=IaBeLJ

## LIST OF SUPPLEMENTARY TABLES

**Supplementary Table S1.** Read alignments in the genomes of *E. coli*, *A. baumannii*, and *P. aeruginos*a.

“S1A” = Supplementary Table S1A. Read alignment statistics for population samples.
“S1B” = Supplementary Table S1B. Read alignment statistics for clonal samples.
“S1C” = Supplementary Table S1C. Genes comprising a mapped large deletion in *A. baumannii ATCC 17978 chromosome*

**Supplementary Table S2.** Observed sequence variants (mutations, small indels, and IS elements insertions in the population samples collected and analyzed from five evolutionary runs.

S2A = Supplementary Table S2A. Sequence variants (nonsynonymous) and their dynamics in evolving populations of *E. coli* BW25113 (run CEC-2)
S2B = Supplementary Table S2B. Sequence variants (nonsynonymous) and their dynamics in evolving populations of *E. coli* BW25113 (run CEC-4)
S2C = Supplementary Table S2C. Sequence variants (nonsynonymous) and their dynamics in evolving populations of *A. baumannii* 17978 (run CAB-1)
S2D= Supplementary Table S2D. Sequence variants (nonsynonymous) and their dynamics in evolving populations of P. aeruginosa (run PAC-1)
S2E= Supplementary Table S2E. Sequence variants (nonsynonymous) and their dynamics in evolving populations of *P. aeruginosa* (run PAC-2)

**Supplementary Table S3.** Observed acquired sequence variants and MIC values in selected clones.

S3A= Supplementary Table S3A. Sequence variants and MIC values in selected clones from the two evolutionary runs of *E. coli* BW25113 (CEC-2 and CEC-4)
S3B = Supplementary Table S3B. Sequence variants and MIC values in selected clones from the evolutionary run of *A. baumannii* ATCC17978 (CAB)
S3C =Supplementary Table S3C. Sequence variants (from directed PCR analysis) and MIC values in selected clones from the evolutionary run of P. aeruginosa ATCC27853 (PAC-1)

## LEGENDS FOR SUPPLEMENTARY FIGURES

**Supplementary Figure S1. Morbidostat-based experimental evolution workflow.** The initial unevolved culture (1) is plated on the agar Petri dish (2) to make individual colonies. The colonies are collected (3) and used to make 6 inoculates (4) – one colony for each reactor for morbidostat cultivation (5). From each reactor samples are taken roughly once in 24 hours (6). To observe population dynamics (8) we perform WGS with high coverage for each sample (7). The population dynamics is used to choose optimal number of colonies from plated evolved samples (9). For each colony we do WGS (10, 11) and MIC tests (12) that reveals genotype-phenotype association.

**Supplementary Figure S2.** OD profiles (black line) and calculated drug concentration profiles (red line) obtained in the course of experimental evolution of ciprofloxacin resistance in (**A**) *Escherichia coli* BW25113, CEC-2 run; (**B**) *Escherichia coli* BW25113, CEC-4 run; (**C**) *Acinetobacter baumannii* ATCC17978, CAB-1 run; (**D**) *Pseudomonas aeruginosa* ATCC27853, PAC-1 run; (**E**) *Pseudomonas aeruginosa* ATCC27853, PAC-2 run. The right axis shows CIP concentration (xMIC) as fold-change relative to MIC value of respective unevolved strains. Sequenced samples marked with arrows.

**Supplementary Figure 3. Computational pipeline for a primary analysis of population sequencing data.** Data shown in black hexagons. Processes and software shown in rectangles and rounded rectangles respectively. Frames indicate aims of the parts of the analysis.

**Supplementary Figure S4.** Population dynamics of experimental evolution of ciprofloxacin resistance in (**A**) *Escherichia coli* BW25113, CEC-2 run; (**B**) *Escherichia coli* BW25113, CEC-4 run; (**C**) *Acinetobacter baumannii* ATCC17978, CAB-1 run; (**D**) *Pseudomonas aeruginosa* ATCC27853, PAC-1 run; (**E**) *Pseudomonas aeruginosa* ATCC27853, PAC-2 run. For each reactor in every morbidostat run, the frequency of major mutations (reaching ≥5%) in evolving bacterial populations is shown as a function of time.

## References

1. Torok E, Moran E, Cooke F. 2010. Oxford Handbook of Infectious Diseases and Microbiology doi:10.1016/j.trstmh.2009.11.002. Oxford University Press, Oxford.

2. Meesters K, Mauel R, Dhont E, Walle JV, De Bruyne P. 2018. Systemic fluoroquinolone prescriptions for hospitalized children in Belgium, results of a multicenter retrospective drug utilization study. BMC Infect Dis 18:89.

3. Pitout JD, Chan WW, Church DL. 2016. Tackling antimicrobial resistance in lower urinary tract infections: treatment options. Expert Rev Anti Infect Ther 14:621–32.

4. Chen CR, Malik M, Snyder M, Drlica K. 1996. DNA gyrase and topoisomerase IV on the bacterial chromosome: quinolone-induced DNA cleavage. J Mol Biol 258:627–37.

5. Drlica K, Malik M, Kerns RJ, Zhao X. 2008. Quinolone-mediated bacterial death. Antimicrob Agents Chemother 52:385–92.

6. Fabrega A, Madurga S, Giralt E, Vila J. 2009. Mechanism of action of and resistance to quinolones. Microb Biotechnol 2:40–61.

7. Hooper DC. 1999. Mode of action of fluoroquinolones. Drugs 58 Suppl 2:6–10.

8. Hooper DC. 2001. Mechanisms of action of antimicrobials: focus on fluoroquinolones. Clin Infect Dis 32 Suppl 1:S9–S15.

9. Gullberg E, Cao S, Berg OG, Ilback C, Sandegren L, Hughes D, Andersson DI. 2011. Selection of resistant bacteria at very low antibiotic concentrations. PLoS Pathog 7:e1002158.

10. Huseby DL, Pietsch F, Brandis G, Garoff L, Tegehall A, Hughes D. 2017. Mutation Supply and Relative Fitness Shape the Genotypes of Ciprofloxacin-Resistant Escherichia coli. Mol Biol Evol 34:1029–1039.

11. Komp Lindgren P, Karlsson A, Hughes D. 2003. Mutation rate and evolution of fluoroquinolone resistance in Escherichia coli isolates from patients with urinary tract infections. Antimicrobial agents and chemotherapy 47:3222–3232.

12. Ruiz J. 2003. Mechanisms of resistance to quinolones: target alterations, decreased accumulation and DNA gyrase protection. J Antimicrob Chemother 51:1109–17.

13. Yoshida H, Bogaki M, Nakamura M, Nakamura S. 1990. Quinolone resistance-determining region in the DNA gyrase gyrA gene of Escherichia coli. Antimicrob Agents Chemother 34:1271–2.

14. Alekshun MN, Levy SB. 1997. Regulation of chromosomally mediated multiple antibiotic resistance: the mar regulon. Antimicrob Agents Chemother 41:2067–75.

15. Cohen SP, McMurry LM, Hooper DC, Wolfson JS, Levy SB. 1989. Cross-resistance to fluoroquinolones in multiple-antibiotic-resistant (Mar) Escherichia coli selected by tetracycline or chloramphenicol: decreased drug accumulation associated with membrane changes in addition to OmpF reduction. Antimicrob Agents Chemother 33:1318–25.

16. Hooper DC, Wolfson JS, Bozza MA, Ng EY. 1992. Genetics and Regulation of Outer Membrane Protein Expression by Quinolone Resistance Loci nfxB, nfxcC, and cfxB. Antimicrobial Agents and Chemotherapy 36:1151–1154.

17. Okusu H, Ma D, Nikaido H. 1996. AcrAB efflux pump plays a major role in the antibiotic resistance phenotype of Escherichia coli multiple-antibiotic-resistance (Mar) mutants. J Bacteriol 178:306–8.

18. Wang H, Dzink-Fox JL, Chen M, Levy SB. 2001. Genetic characterization of highly fluoroquinolone-resistant clinical Escherichia coli strains from China: role of acrR mutations. Antimicrob Agents Chemother 45:1515–21.

19. Lee CR, Lee JH, Park M, Park KS, Bae IK, Kim YB, Cha CJ, Jeong BC, Lee SH. 2017. Biology of Acinetobacter baumannii: Pathogenesis, Antibiotic Resistance Mechanisms, and Prospective Treatment Options. Front Cell Infect Microbiol 7:55.

20. McConnell MJ, Actis L, Pachon J. 2013. Acinetobacter baumannii: human infections, factors contributing to pathogenesis and animal models. FEMS Microbiol Rev 37:130–55.

21. Park S, Lee KM, Yoo YS, Yoo JS, Yoo JI, Kim HS, Lee YS, Chung GT. 2011. Alterations of gyrA, gyrB, and parC and Activity of Efflux Pump in Fluoroquinolone-resistant Acinetobacter baumannii. Osong Public Health Res Perspect 2:164–70.

22. Rehman A, Patrick WM, Lamont IL. 2019. Mechanisms of ciprofloxacin resistance in Pseudomonas aeruginosa: new approaches to an old problem. J Med Microbiol 68:1–10.

23. Garoff L, Pietsch F, Huseby DL, Lilja T, Brandis G, Hughes D. 2020. Population Bottlenecks Strongly Influence the Evolutionary Trajectory to Fluoroquinolone Resistance in Escherichia coli. Mol Biol Evol 37:1637–1646.

24. Komp Lindgren P, Marcusson LL, Sandvang D, Frimodt-Moller N, Hughes D. 2005. Biological cost of single and multiple norfloxacin resistance mutations in Escherichia coli implicated in urinary tract infections. Antimicrob Agents Chemother 49:2343–51.

25. Zhao X, Drlica K. 2001. Restricting the selection of antibiotic-resistant mutants: a general strategy derived from fluoroquinolone studies. Clin Infect Dis 33 Suppl 3:S147–56.

26. Toprak E, Veres A, Michel JB, Chait R, Hartl DL, Kishony R. 2011. Evolutionary paths to antibiotic resistance under dynamically sustained drug selection. Nat Genet 44:101–5.

27. Toprak E, Veres A, Yildiz S, Pedraza JM, Chait R, Paulsson J, Kishony R. 2013. Building a morbidostat: an automated continuous-culture device for studying bacterial drug resistance under dynamically sustained drug inhibition. Nat Protoc 8:555–67.

28. Leyn SA, Zlamal JE, Kurnasov OV, Li X, Elane M, Myjak L, Godzik M, de Crecy A, Garcia FA, Ebeling M, Osterman AL. 2021. Experimental evolution in morbidostat reveals converging genomic trajectories on the path to triclosan resistance. Microb Genom In Press.

29. Andrews S. 2010. FastQC. A quality control tool for high throughput sequence data, Babraham Institute, https://www.bioinformatics.babraham.ac.uk/projects/fastqc/.

30. Bolger AM, Lohse M, Usadel B. 2014. Trimmomatic: a flexible trimmer for Illumina sequence data. Bioinformatics 30:2114–20.

31. Li H, Durbin R. 2009. Fast and accurate short read alignment with Burrows-Wheeler transform. Bioinformatics 25:1754–60.

32. Wilm A, Aw PP, Bertrand D, Yeo GH, Ong SH, Wong CH, Khor CC, Petric R, Hibberd ML, Nagarajan N. 2012. LoFreq: a sequence-quality aware, ultra-sensitive variant caller for uncovering cell-population heterogeneity from high-throughput sequencing datasets. Nucleic Acids Res 40:11189–201.

33. DePristo MA, Banks E, Poplin R, Garimella KV, Maguire JR, Hartl C, Philippakis AA, del Angel G, Rivas MA, Hanna M, McKenna A, Fennell TJ, Kernytsky AM, Sivachenko AY, Cibulskis K, Gabriel SB, Altshuler D, Daly MJ. 2011. A framework for variation discovery and genotyping using next-generation DNA sequencing data. Nat Genet 43:491–8.

34. Siguier P, Perochon J, Lestrade L, Mahillon J, Chandler M. 2006. ISfinder: the reference centre for bacterial insertion sequences. Nucleic Acids Res 34:D32–6.

35. Cingolani P, Platts A, Wang le L, Coon M, Nguyen T, Wang L, Land SJ, Lu X, Ruden DM. 2012. A program for annotating and predicting the effects of single nucleotide polymorphisms, SnpEff: SNPs in the genome of Drosophila melanogaster strain w1118; iso-2; iso-3. Fly (Austin) 6:80–92.

36. Li H. 2011. A statistical framework for SNP calling, mutation discovery, association mapping and population genetical parameter estimation from sequencing data. Bioinformatics 27:2987–93.

37. Deatherage DE, Barrick JE. 2014. Identification of mutations in laboratory-evolved microbes from next-generation sequencing data using breseq. Methods Mol Biol 1151:165–88.

38. Brynildsrud O, Snipen LG, Bohlin J. 2015. CNOGpro: detection and quantification of CNVs in prokaryotic whole-genome sequencing data. Bioinformatics 31:1708–15.

39. Kurtz S, Phillippy A, Delcher AL, Smoot M, Shumway M, Antonescu C, Salzberg SL. 2004. Versatile and open software for comparing large genomes. Genome Biol 5:R12.

40. Bankevich A, Nurk S, Antipov D, Gurevich AA, Dvorkin M, Kulikov AS, Lesin VM, Nikolenko SI, Pham S, Prjibelski AD, Pyshkin AV, Sirotkin AV, Vyahhi N, Tesler G, Alekseyev MA, Pevzner PA. 2012. SPAdes: a new genome assembly algorithm and its applications to single-cell sequencing. J Comput Biol 19:455–77.

41. Brettin T, Davis JJ, Disz T, Edwards RA, Gerdes S, Olsen GJ, Olson R, Overbeek R, Parrello B, Pusch GD, Shukla M, Thomason JA, 3rd, Stevens R, Vonstein V, Wattam AR, Xia F. 2015. RASTtk: a modular and extensible implementation of the RAST algorithm for building custom annotation pipelines and annotating batches of genomes. Sci Rep 5:8365.

42. EUCAST. 2000. EUCAST Definitive Document E.DEF 3.1, June 2000: Determination of minimum inhibitory concentrations (MICs) of antibacterial agents by agar dilution. Clin Microbiol Infect 6:509–15.

43. Davis JJ, Wattam AR, Aziz RK, Brettin T, Butler R, Butler RM, Chlenski P, Conrad N, Dickerman A, Dietrich EM, Gabbard JL, Gerdes S, Guard A, Kenyon RW, Machi D, Mao C, Murphy-Olson D, Nguyen M, Nordberg EK, Olsen GJ, Olson RD, Overbeek JC, Overbeek R, Parrello B, Pusch GD, Shukla M, Thomas C, VanOeffelen M, Vonstein V, Warren AS, Xia F, Xie D, Yoo H, Stevens R. 2020. The PATRIC Bioinformatics Resource Center: expanding data and analysis capabilities. Nucleic Acids Res 48:D606–D612.

44. Marcusson LL, Frimodt-Moller N, Hughes D. 2009. Interplay in the selection of fluoroquinolone resistance and bacterial fitness. PLoS Pathog 5:e1000541.

45. Hopkins KL, Davies RH, Threlfall EJ. 2005. Mechanisms of quinolone resistance in Escherichia coli and Salmonella: recent developments. Int J Antimicrob Agents 25:358–73.

46. Minnick MF, Wilson ZR, Smitherman LS, Samuels DS. 2003. gyrA mutations in ciprofloxacin-resistant Bartonella bacilliformis strains obtained in vitro. Antimicrob Agents Chemother 47:383–6.

47. Bagel S, Hullen V, Wiedemann B, Heisig P. 1999. Impact of gyrA and parC mutations on quinolone resistance, doubling time, and supercoiling degree of Escherichia coli. Antimicrob Agents Chemother 43:868–75.

48. Bhatnagar K, Wong A. 2019. The mutational landscape of quinolone resistance in Escherichia coli. PLoS One 14:e0224650.

49. Komp Lindgren P, Karlsson A, Hughes D. 2003. Mutation rate and evolution of fluoroquinolone resistance in Escherichia coli isolates from patients with urinary tract infections. Antimicrob Agents Chemother 47:3222–32.

50. Vinue L, Corcoran MA, Hooper DC, Jacoby GA. 2015. Mutations That Enhance the Ciprofloxacin Resistance of Escherichia coli with qnrA1. Antimicrob Agents Chemother 60:1537–45.

51. Maneewannakul K, Levy SB. 1996. Identification for mar mutants among quinolone-resistant clinical isolates of Escherichia coli. Antimicrob Agents Chemother 40:1695–8.

52. Linde HJ, Notka F, Metz M, Kochanowski B, Heisig P, Lehn N. 2000. In Vivo Increase in Resistance to Ciprofloxacin in Escherichia coli Associated with Deletion of the C-Terminal Part of MarR. Antimicrobial Agents and Chemotherapy 44:1865–1868.

53. Alekshun MN, Levy SB. 1999. The mar regulon: multiple resistance to antibiotics and other toxic chemicals. Trends Microbiol 7:410–3.

54. Zhu R, Hao Z, Lou H, Song Y, Zhao J, Chen Y, Zhu J, Chen PR. 2017. Structural characterization of the DNA-binding mechanism underlying the copper(II)-sensing MarR transcriptional regulator. J Biol Inorg Chem 22:685–693.

55. Praski Alzrigat L, Huseby DL, Brandis G, Hughes D. 2017. Fitness cost constrains the spectrum of marR mutations in ciprofloxacin-resistant Escherichia coli. J Antimicrob Chemother 72:3016–3024.

56. Weber BS, Ly PM, Irwin JN, Pukatzki S, Feldman MF. 2015. A multidrug resistance plasmid contains the molecular switch for type VI secretion in Acinetobacter baumannii. Proc Natl Acad Sci U S A 112:9442–7.

57. Higuchi S, Onodera Y, Chiba M, Hoshino K, Gotoh N. 2013. Potent in vitro antibacterial activity of DS-8587, a novel broad-spectrum quinolone, against Acinetobacter baumannii. Antimicrob Agents Chemother 57:1978–81.

58. Gerson S, Nowak J, Zander E, Ertel J, Wen Y, Krut O, Seifert H, Higgins PG. 2018. Diversity of mutations in regulatory genes of resistance-nodulation-cell division efflux pumps in association with tigecycline resistance in Acinetobacter baumannii. J Antimicrob Chemother 73:1501–1508.

59. Marchand I, Damier-Piolle L, Courvalin P, Lambert T. 2004. Expression of the RND-type efflux pump AdeABC in Acinetobacter baumannii is regulated by the AdeRS two-component system. Antimicrob Agents Chemother 48:3298–304.

60. Xu C, Bilya SR, Xu W. 2019. adeABC efflux gene in Acinetobacter baumannii. New Microbes New Infect 30:100549.

61. Damier-Piolle L, Magnet S, Bremont S, Lambert T, Courvalin P. 2008. AdeIJK, a resistance-nodulation-cell division pump effluxing multiple antibiotics in Acinetobacter baumannii. Antimicrob Agents Chemother 52:557–62.

62. Rosenfeld N, Bouchier C, Courvalin P, Perichon B. 2012. Expression of the resistance-nodulation-cell division pump AdeIJK in Acinetobacter baumannii is regulated by AdeN, a TetR-type regulator. Antimicrob Agents Chemother 56:2504–10.

63. Sugawara E, Nikaido H. 2014. Properties of AdeABC and AdeIJK efflux systems of Acinetobacter baumannii compared with those of the AcrAB-TolC system of Escherichia coli. Antimicrob Agents Chemother 58:7250–7.

64. Coyne S, Courvalin P, Perichon B. 2011. Efflux-mediated antibiotic resistance in Acinetobacter spp. Antimicrob Agents Chemother 55:947–53.

65. Lee JK, Lee YS, Park YK, Kim BS. 2005. Mutations in the gyrA and parC genes in ciprofloxacin-resistant clinical isolates of Acinetobacter baumannii in Korea. Microbiol Immunol 49:647–53.

66. Vila J, Ruiz J, Goni P, Jimenez de Anta T. 1997. Quinolone-resistance mutations in the topoisomerase IV parC gene of Acinetobacter baumannii. J Antimicrob Chemother 39:757–62.

67. Qin TT, Kang HQ, Ma P, Li PP, Huang LY, Gu B. 2015. SOS response and its regulation on the fluoroquinolone resistance. Ann Transl Med 3:358.

68. Lee JK, Lee YS, Park YK, Kim BS. 2005. Alterations in the GyrA and GyrB subunits of topoisomerase II and the ParC and ParE subunits of topoisomerase IV in ciprofloxacin-resistant clinical isolates of Pseudomonas aeruginosa. Int J Antimicrob Agents 25:290–5.

69. Yonezawa M, Takahata M, Matsubara N, Watanabe Y, Narita H. 1995. DNA gyrase gyrA mutations in quinolone-resistant clinical isolates of Pseudomonas aeruginosa. Antimicrob Agents Chemother 39:1970–2.

70. Poole K, Gotoh N, Tsujimoto H, Zhao Q, Wada A, Yamasaki T, Neshat S, Yamagishi J, Li XZ, Nishino T. 1996. Overexpression of the mexC-mexD-oprJ efflux operon in nfxB-type multidrug-resistant strains of Pseudomonas aeruginosa. Mol Microbiol 21:713–24.

71. Shiba T, Ishiguro K, Takemoto N, Koibuchi H, Sugimoto K. 1995. Purification and characterization of the Pseudomonas aeruginosa NfxB protein, the negative regulator of the nfxB gene. Journal of Bacteriology 177:5872–5877.

72. Hooper DC, Wolfson JS, Bozza MA, Ng EY. 1992. Genetics and regulation of outer membrane protein expression by quinolone resistance loci nfxB, nfxC, and cfxB. Antimicrob Agents Chemother 36:1151–4.

73. Stickland HG, Davenport PW, Lilley KS, Griffin JL, Welch M. 2010. Mutation of nfxB causes global changes in the physiology and metabolism of Pseudomonas aeruginosa. J Proteome Res 9:2957–67.

74. Purssell A, Poole K. 2013. Functional characterization of the NfxB repressor of the mexCD-oprJ multidrug efflux operon of Pseudomonas aeruginosa. Microbiology (Reading) 159:2058–2073.

75. Kohler T, Epp SF, Curty LK, Pechere JC. 1999. Characterization of MexT, the regulator of the MexE-MexF-OprN multidrug efflux system of Pseudomonas aeruginosa. J Bacteriol 181:6300–5.

76. Sobel ML, Neshat S, Poole K. 2005. Mutations in PA2491 (mexS) promote MexT-dependent mexEF-oprN expression and multidrug resistance in a clinical strain of Pseudomonas aeruginosa. J Bacteriol 187:1246–53.

77. Richardot C, Juarez P, Jeannot K, Patry I, Plesiat P, Llanes C. 2016. Amino Acid Substitutions Account for Most MexS Alterations in Clinical nfxC Mutants of Pseudomonas aeruginosa. Antimicrob Agents Chemother 60:2302–10.

78. Wong A, Kassen R. 2011. Parallel evolution and local differentiation in quinolone resistance in Pseudomonas aeruginosa. Microbiology (Reading) 157:937–944.

79. Ahmed MN, Abdelsamad A, Wassermann T, Porse A, Becker J, Sommer MOA, Hoiby N, Ciofu O. 2020. The evolutionary trajectories of P. aeruginosa in biofilm and planktonic growth modes exposed to ciprofloxacin: beyond selection of antibiotic resistance. NPJ Biofilms Microbiomes 6:28.

80. Ahmed MN, Porse A, Sommer MOA, Hoiby N, Ciofu O. 2018. Evolution of Antibiotic Resistance in Biofilm and Planktonic Pseudomonas aeruginosa Populations Exposed to Subinhibitory Levels of Ciprofloxacin. Antimicrob Agents Chemother 62.

81. Monti MR, Morero NR, Miguel V, Argarana CE. 2013. nfxB as a novel target for analysis of mutation spectra in Pseudomonas aeruginosa. PLoS One 8:e66236.

82. Mena A, Smith EE, Burns JL, Speert DP, Moskowitz SM, Perez JL, Oliver A. 2008. Genetic adaptation of Pseudomonas aeruginosa to the airways of cystic fibrosis patients is catalyzed by hypermutation. Journal of Bacteriology 190:7910–7917.

83. Willmott CJ, Maxwell A. 1993. A single point mutation in the DNA gyrase A protein greatly reduces binding of fluoroquinolones to the gyrase-DNA complex. Antimicrob Agents Chemother 37:126–7.

84. Jacoby GA. 2005. Mechanisms of resistance to quinolones. Clin Infect Dis 41 Suppl 2:S120–6.

85. Gensberg K, Jin YF, Piddock LJ. 1995. A novel gyrB mutation in a fluoroquinolone-resistant clinical isolate of Salmonella typhimurium. FEMS Microbiol Lett 132:57–60.

86. Lascols C, Robert J, Cattoir V, Bebear C, Cavallo JD, Podglajen I, Ploy MC, Bonnet R, Soussy CJ, Cambau E. 2007. Type II topoisomerase mutations in clinical isolates of Enterobacter cloacae and other enterobacterial species harbouring the qnrA gene. Int J Antimicrob Agents 29:402–9.

87. Mouneimne H, Robert J, Jarlier V, Cambau E. 1999. Type II topoisomerase mutations in ciprofloxacin-resistant strains of Pseudomonas aeruginosa. Antimicrob Agents Chemother 43:62–6.

88. Wimalasena S, Pathirana H, Shin GW, De Silva BCJ, Hossain S, Heo GJ. 2019. Characterization of Quinolone-Resistant Determinants in Tribe Proteeae Isolated from Pet Turtles with High Prevalence of qnrD and Novel gyrB Mutations. Microb Drug Resist 25:611–618.

89. Bansal S, Tandon V. 2011. Contribution of mutations in DNA gyrase and topoisomerase IV genes to ciprofloxacin resistance in Escherichia coli clinical isolates. Int J Antimicrob Agents 37:253–5.

90. Vogwill T, MacLean RC. 2015. The genetic basis of the fitness costs of antimicrobial resistance: a meta-analysis approach. Evol Appl 8:284–95.

91. Dosselmann B, Willmann M, Steglich M, Bunk B, Nubel U, Peter S, Neher RA. 2017. Rapid and Consistent Evolution of Colistin Resistance in Extensively Drug-Resistant Pseudomonas aeruginosa during Morbidostat Culture. Antimicrob Agents Chemother 61.

