## Supplementary material for "Shared and Unique Evolutionary Trajectories to Ciprofloxacin Resistance in Gram-negative Bacterial Pathogens": Figure S4

**Supplementary Figure S4.** Population dynamics of experimental evolution of ciprofloxacin resistance in (**A**) *Escherichia coli* BW25113, CEC-2 run; (**B**) *Escherichia coli* BW25113, CEC-4 run; (**C**) *Acinetobacter baumannii* ATCC17978, CAB-1 run; (**D**) *Pseudomonas aeruginosa* ATCC27853, PAC-1 run; (**E**) *Pseudomonas aeruginosa* ATCC27853, PAC-2 run. For each reactor in every morbidostat run, the frequency of major mutations (reaching ≥5%) in evolving bacterial populations is shown as a function of time.

A

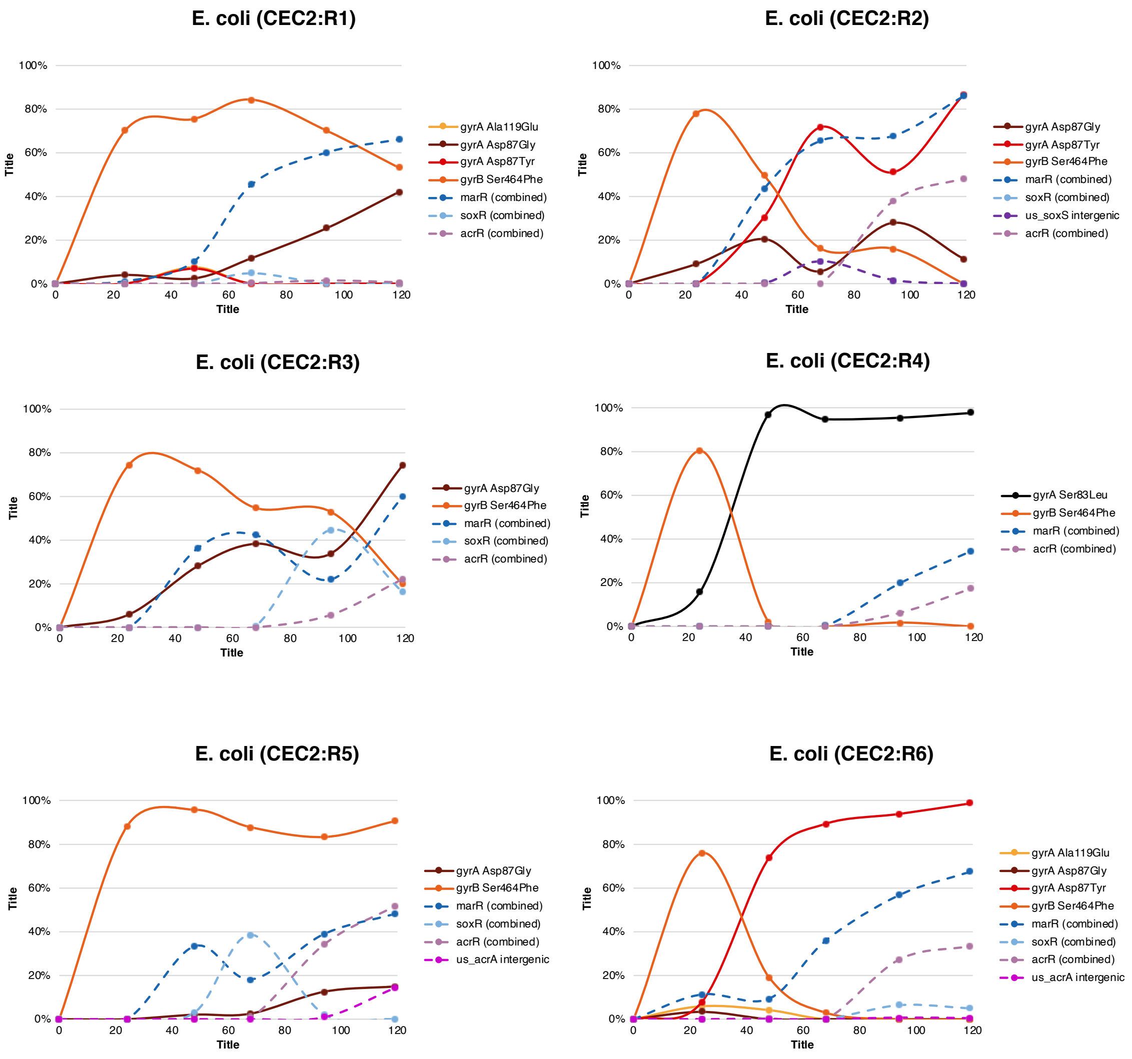

B

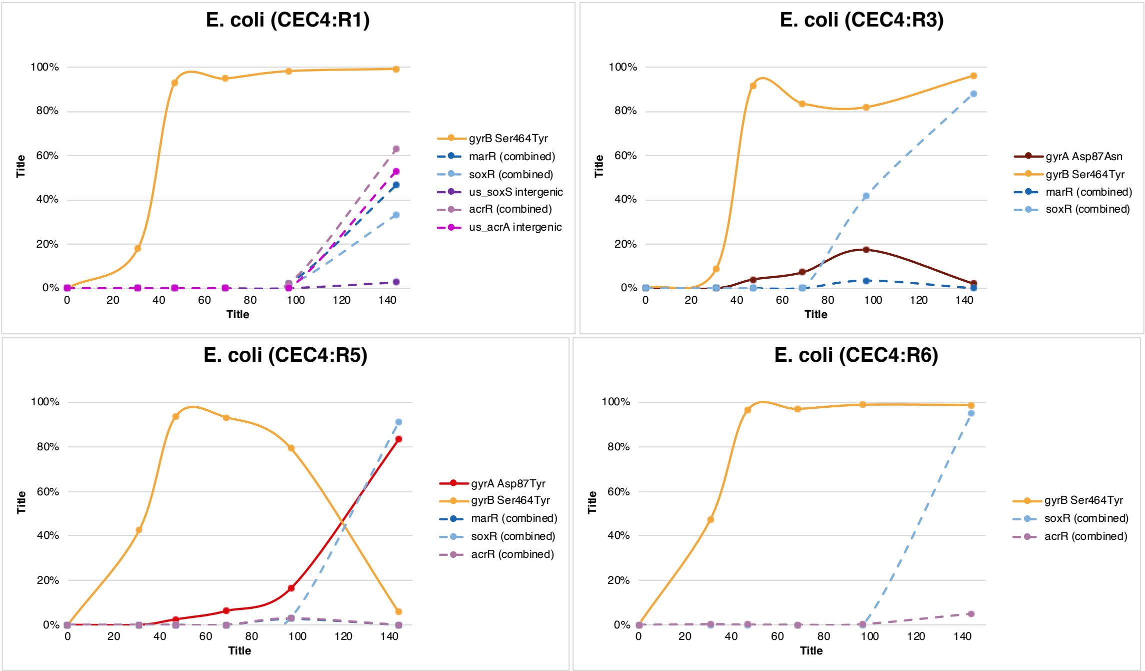
E. coli (CEC4:R1)
100%
80%
60%
40%
20%
0%
Title
0
20
40
60
80
100
120
140
Title
gyrB Ser464Tyr
marR (combined)
soxR (combined)
us_soxS intergenic
acrR (combined)
us_acrA intergenic
E. coli (CEC4:R3)
gyrA Asp87Asn
gyrB Ser464Tyr
marR (combined)
soxR (combined)
E. coli (CEC4:R5)
gyrA Asp87Tyr
gyrB Ser464Tyr
marR (combined)
soxR (combined)
acrR (combined)
E. coli (CEC4:R6)
gyrB Ser464Tyr
soxR (combined)
acrR (combined)

C

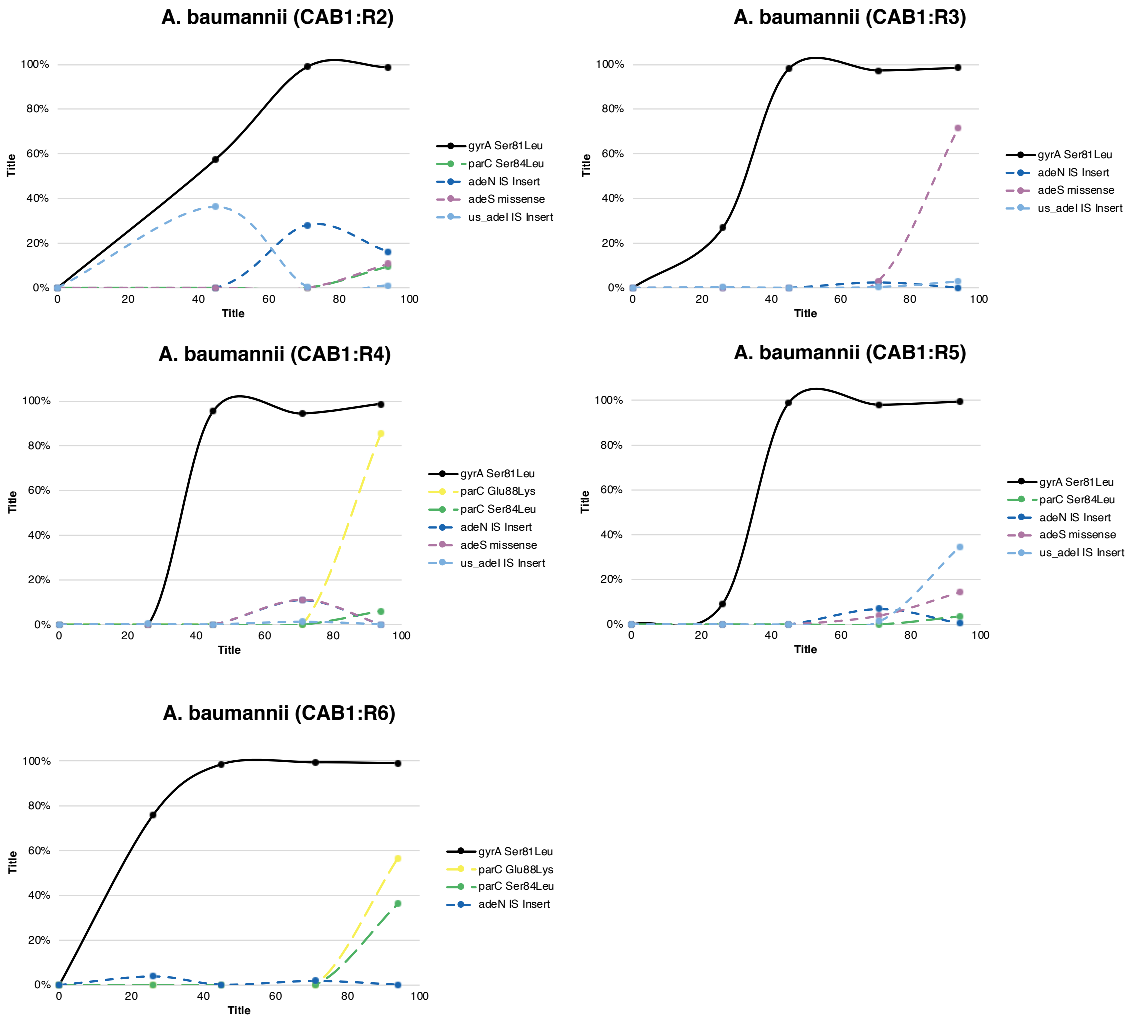

A. baumannii (CAB1:R2)
100%
80%
60%
40%
20%
0%
0
20
40
60
80
100
Title
Title
gyrA Ser81Leu
parC Ser84Leu
adeN IS Insert
adeS missense
us_adeI IS Insert
A. baumannii (CAB1:R3)
gyrA Ser81Leu
adeN IS Insert
adeS missense
us_adeI IS Insert
A. baumannii (CAB1:R4)
gyrA Ser81Leu
parC Glu88Lys
parC Ser84Leu
adeN IS Insert
adeS missense
us_adeI IS Insert
A. baumannii (CAB1:R5)
gyrA Ser81Leu
parC Ser84Leu
adeN IS Insert
adeS missense
us_adeI IS Insert
A. baumannii (CAB1:R6)
gyrA Ser81Leu
parC Glu88Lys
parC Ser84Leu
adeN IS Insert

D

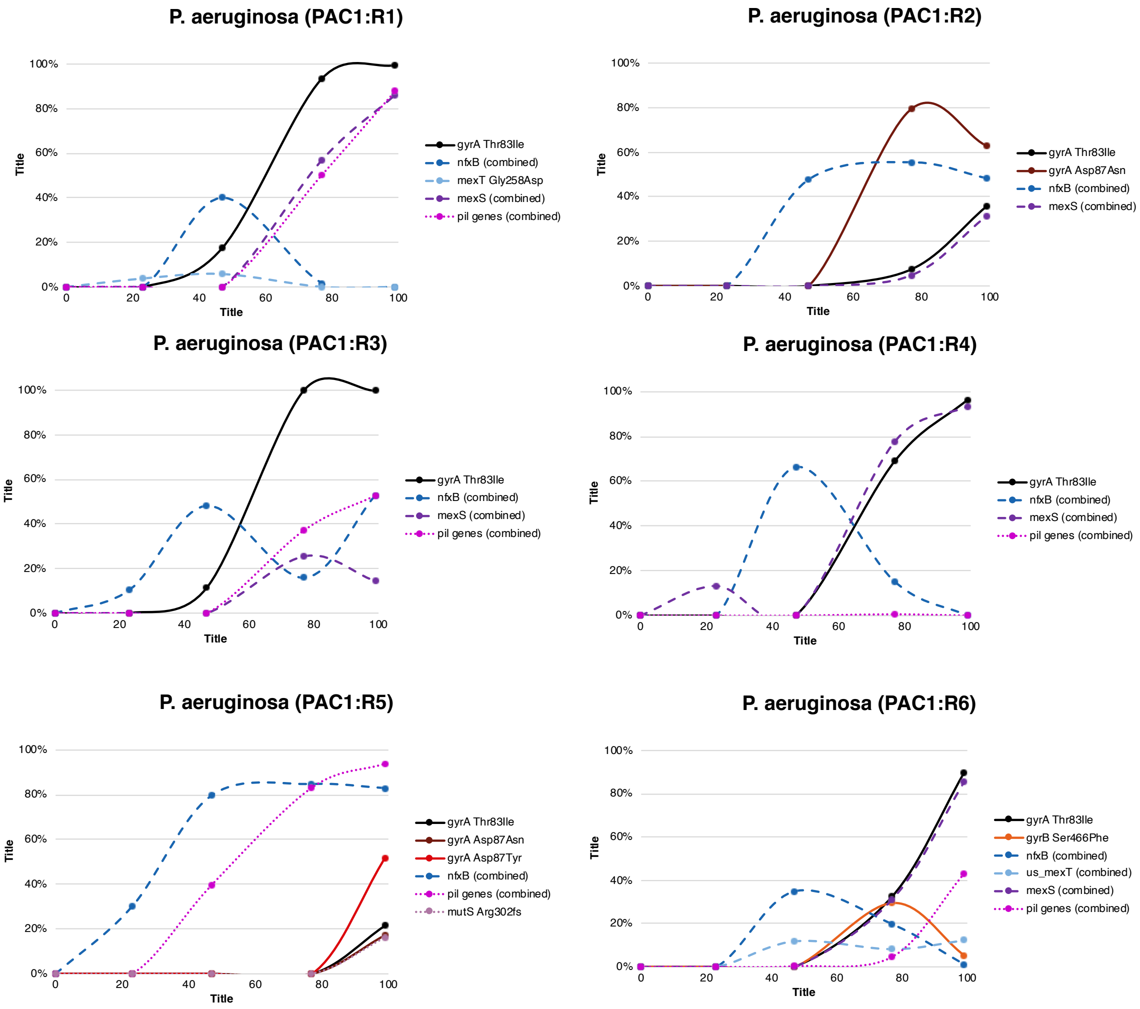
P. aeruginosa (PAC1:R1)
100%
80%
60%
40%
20%
0%
0
20
40
60
80
100
Title
Title
gyrA Thr83Ile
nfxB (combined)
mexT Gly258Asp
mexS (combined)
pil genes (combined)
P. aeruginosa (PAC1:R2)
gyrA Thr83Ile
gyrA Asp87Asn
nfxB (combined)
mexS (combined)
P. aeruginosa (PAC1:R3)
gyrA Thr83Ile
nfxB (combined)
mexS (combined)
pil genes (combined)
P. aeruginosa (PAC1:R4)
gyrA Thr83Ile
nfxB (combined)
mexS (combined)
pil genes (combined)
P. aeruginosa (PAC1:R5)
gyrA Thr83Ile
gyrA Asp87Asn
gyrA Asp87Tyr
nfxB (combined)
pil genes (combined)
mutS Arg302fs
P. aeruginosa (PAC1:R6)
gyrA Thr83Ile
gyrB Ser466Phe
nfxB (combined)
us_mexT (combined)
mexS (combined)
pil genes (combined)

E

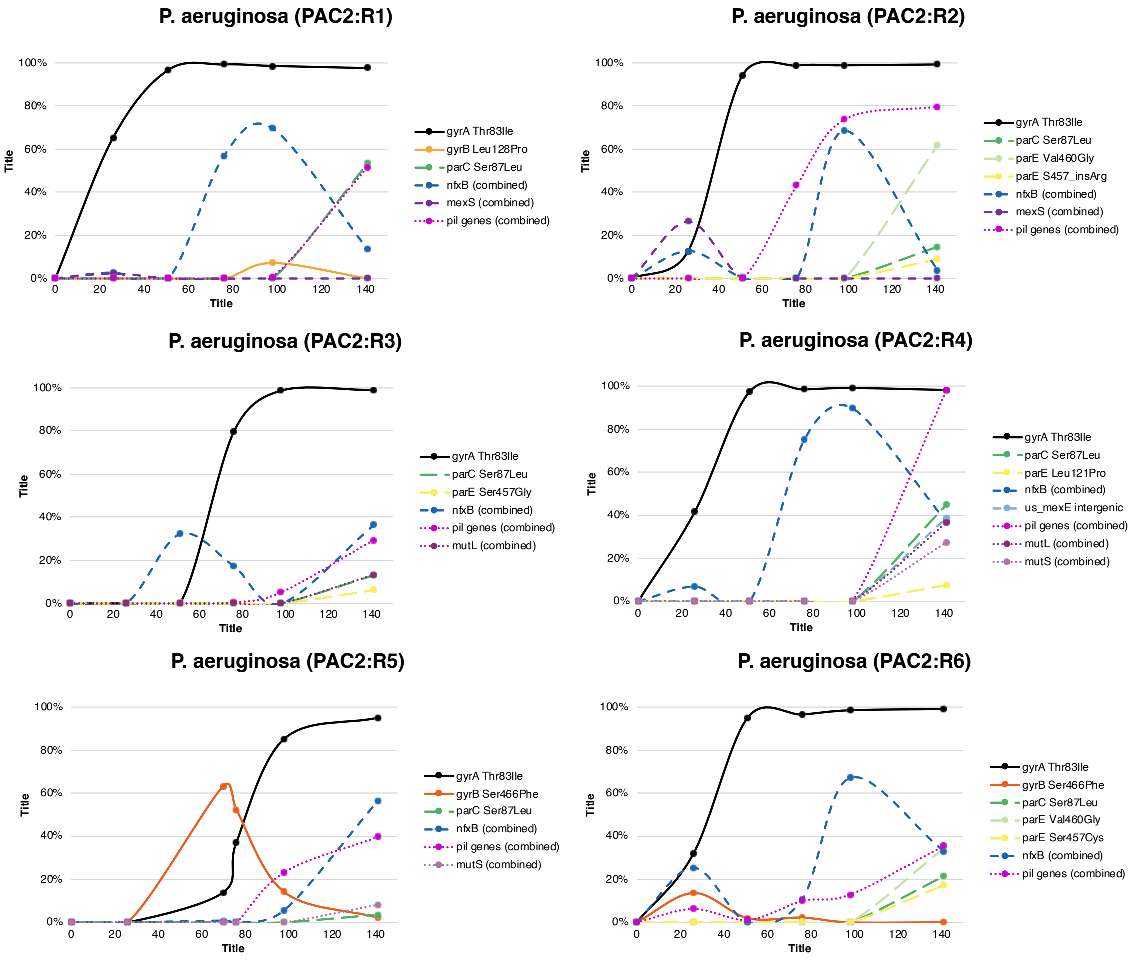
P. aeruginosa (PAC2:R1)
100%
80%
60%
40%
20%
0%
Title
0
20
40
60
80
100
120
140
Title
gyrA Thr83Ile
gyrB Leu128Pro
parC Ser87Leu
nfxB (combined)
mexS (combined)
pil genes (combined)
P. aeruginosa (PAC2:R2)
gyrA Thr83Ile
parC Ser87Leu
parE Val460Gly
parE S457_insArg
nfxB (combined)
mexS (combined)
pil genes (combined)
P. aeruginosa (PAC2:R3)
gyrA Thr83Ile
parC Ser87Leu
parE Ser457Gly
nfxB (combined)
pil genes (combined)
mutL (combined)
P. aeruginosa (PAC2:R4)
gyrA Thr83Ile
parC Ser87Leu
parE Leu121Pro
nfxB (combined)
us_mexE intergenic
pil genes (combined)
mutL (combined)
mutS (combined)
P. aeruginosa (PAC2:R5)
gyrA Thr83Ile
gyrB Ser466Phe
parC Ser87Leu
nfxB (combined)
pil genes (combined)
mutS (combined)
P. aeruginosa (PAC2:R6)
gyrA Thr83Ile
gyrB Ser466Phe
parC Ser87Leu
parE Val460Gly
parE Ser457Cys
nfxB (combined)
pil genes (combined)
