## Supplementary material for "Shared and Unique Evolutionary Trajectories to Ciprofloxacin Resistance in Gram-negative Bacterial Pathogens": Figure S3

**Supplementary Figure 3. Computational pipeline for a primary analysis of population sequencing data.** Data shown in black hexagons. Processes and software shown in rectangles and rounded rectangles respectively. Frames indicate aims of the parts of the analysis.

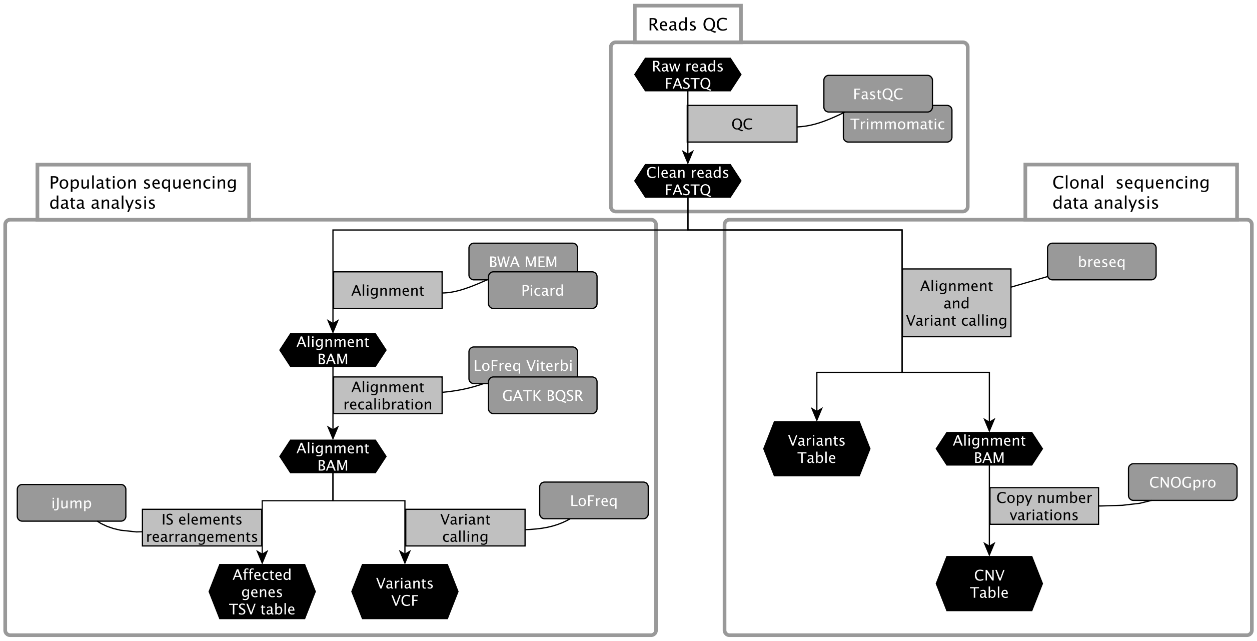
