## Supplementary material for "Shared and Unique Evolutionary Trajectories to Ciprofloxacin Resistance in Gram-negative Bacterial Pathogens": Figure S2

**Supplementary Figure S2.** OD profiles (black line) and calculated drug concentration profiles (red line) obtained in the course of experimental evolution of ciprofloxacin resistance in (**A**) *Escherichia coli* BW25113, CEC-2 run; (**B**) *Escherichia coli* BW25113, CEC-4 run; (**C**) *Acinetobacter baumannii* ATCC17978, CAB-1 run; (**D**) *Pseudomonas aeruginosa* ATCC27853, PAC-1 run; (**E**) *Pseudomonas aeruginosa* ATCC27853, PAC-2 run. The right axis shows CIP concentration (xMIC) as fold-change relative to MIC value of respective unevolved strains. Sequenced samples marked with arrows.

A

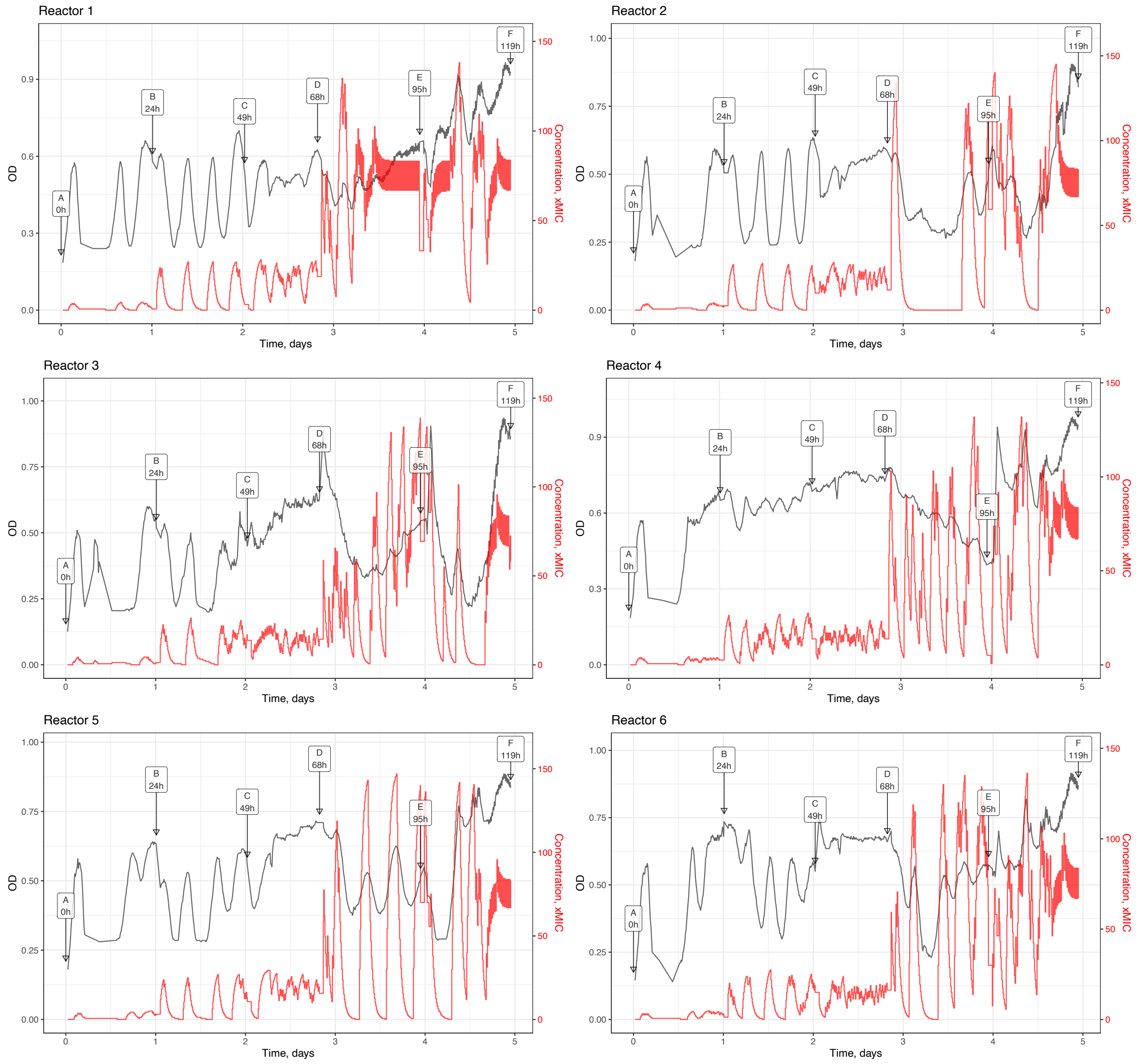

B

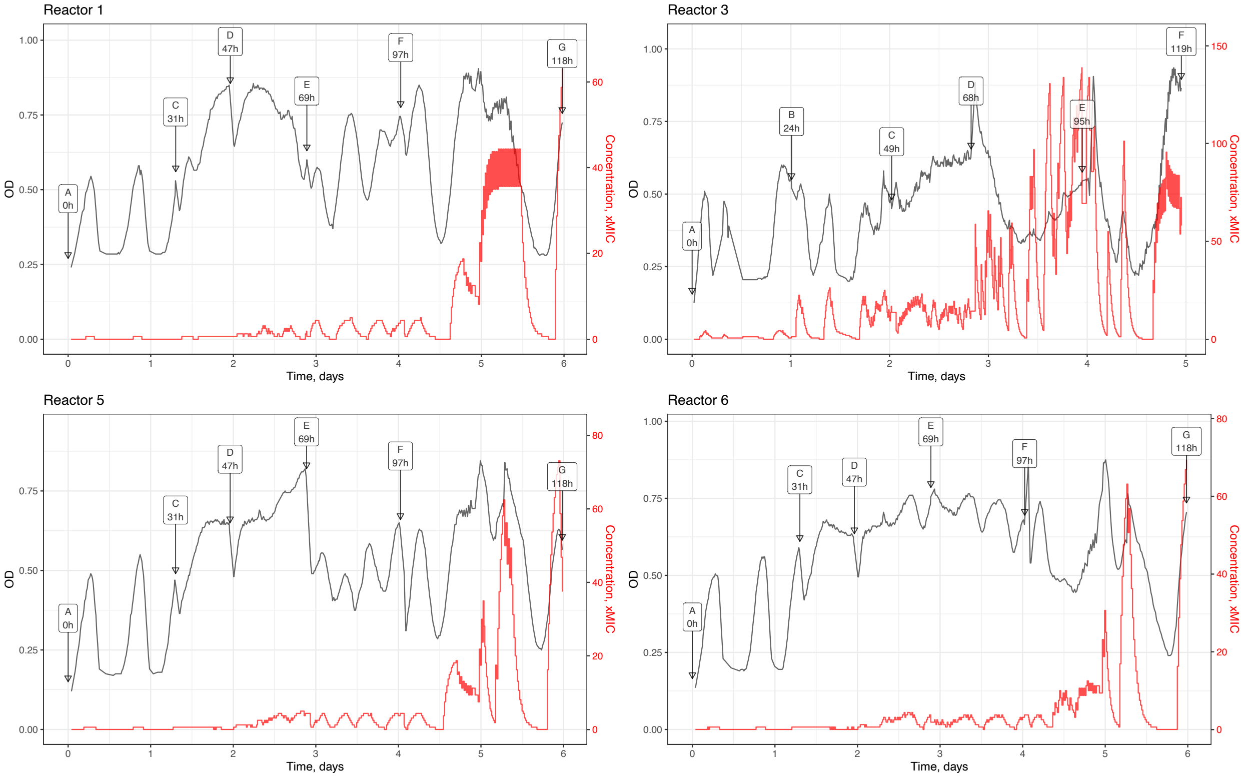
Reactor 1
Reactor 3
Reactor 5
Reactor 6
OD
Concentration, xMIC
Time, days
A 0h
B 24h
C 31h
C 49h
D 47h
D 68h
E 69h
E 95h
F 97h
F 119h
G 118h

C

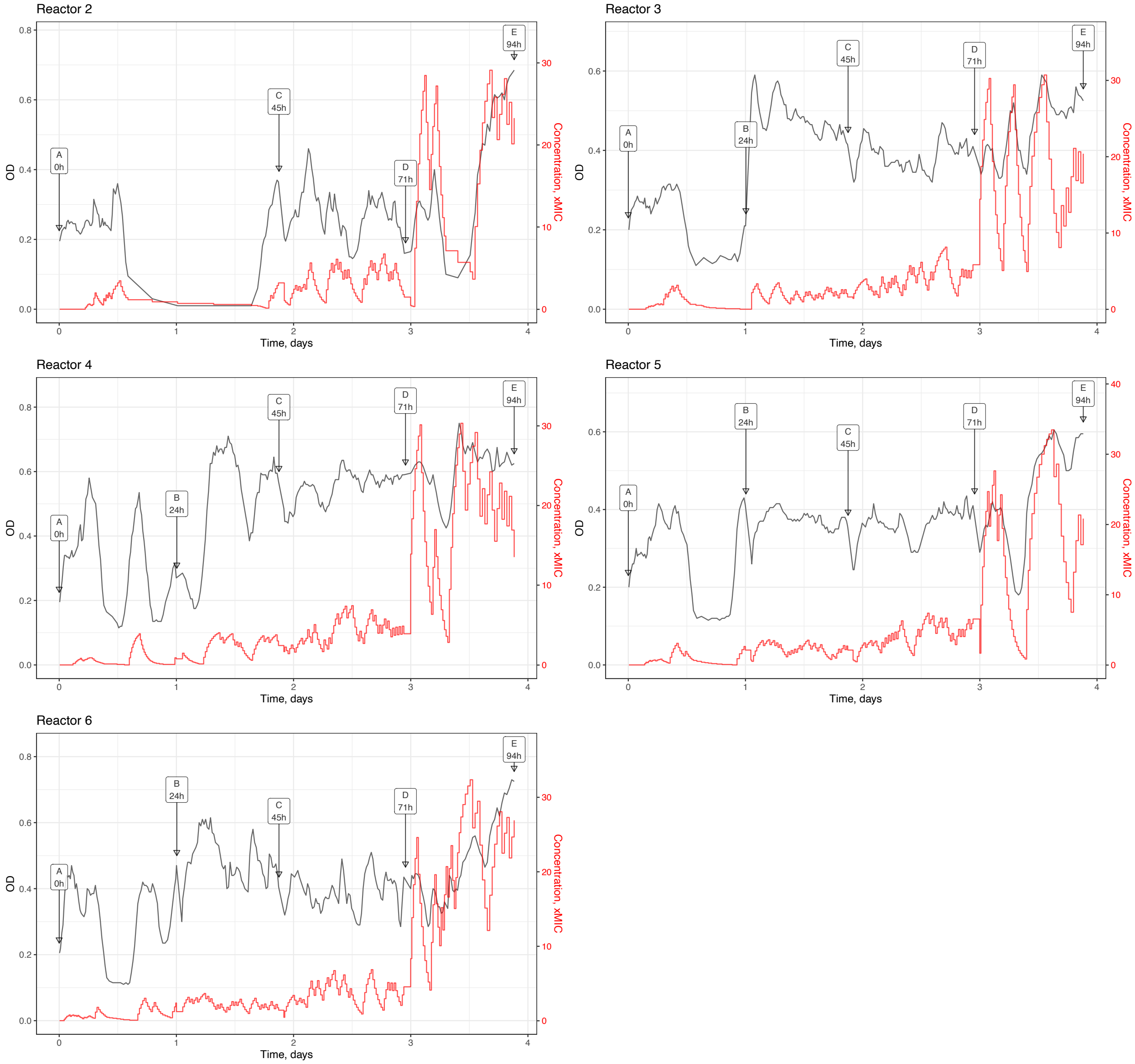
Reactor 2
Reactor 3
Reactor 4
Reactor 5
Reactor 6
OD
Concentration, xMIC
Time, days
A
0h
B
24h
C
45h
D
71h
E
94h
0.0
0.2
0.4
0.6
0.8
0
10
20
30
40
1
2
3
4

D

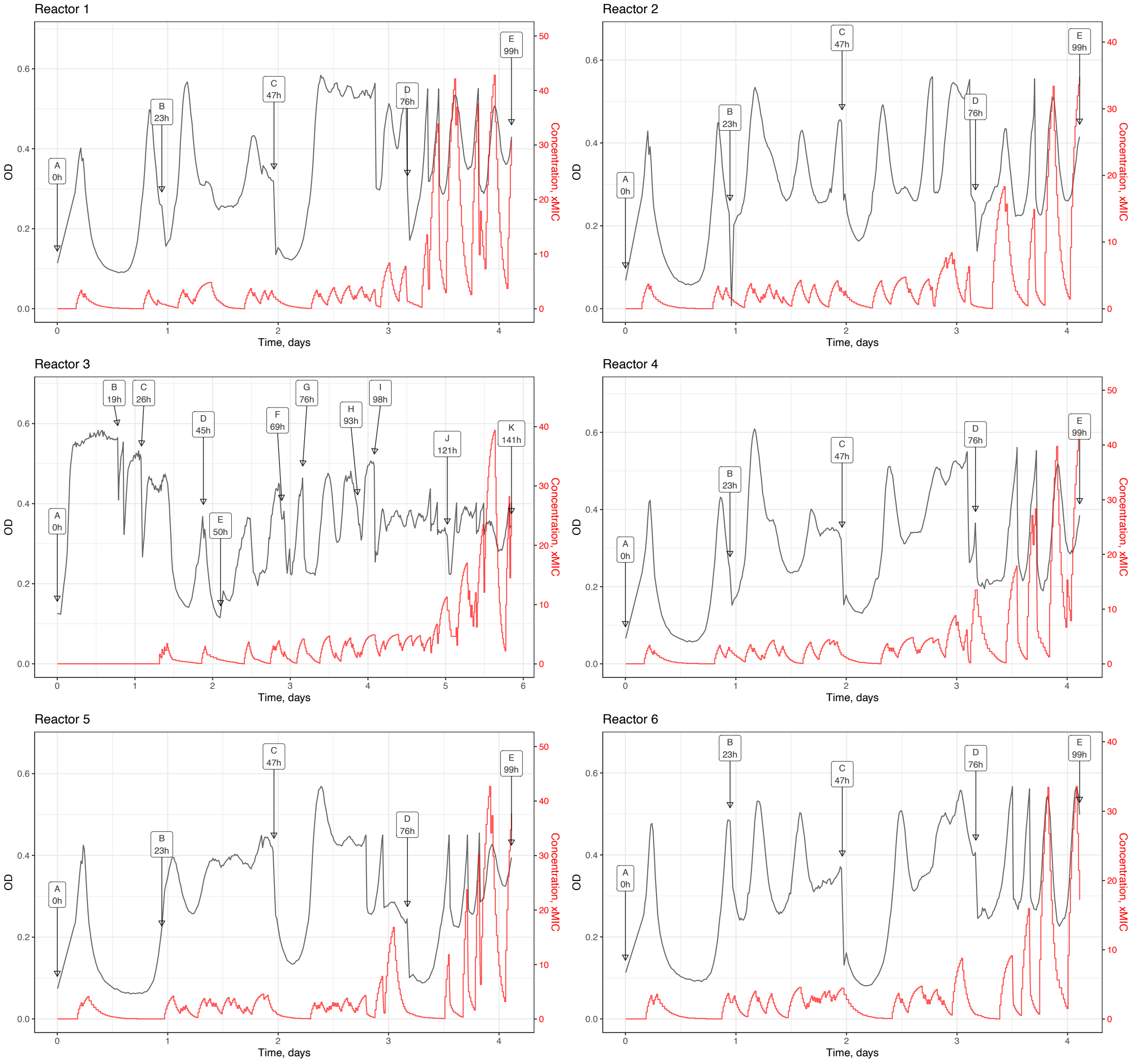

Reactor 1
A 0h
B 23h
C 47h
D 76h
E 99h
OD
Concentration, xMIC
Time, days
Reactor 2
A 0h
B 23h
C 47h
D 76h
E 99h
OD
Concentration, xMIC
Time, days
Reactor 3
A 0h
B 19h
C 26h
D 45h
E 50h
F 69h
G 76h
H 93h
I 98h
J 121h
K 141h
OD
Concentration, xMIC
Time, days
Reactor 4
A 0h
B 23h
C 47h
D 76h
E 99h
OD
Concentration, xMIC
Time, days
Reactor 5
A 0h
B 23h
C 47h
D 76h
E 99h
OD
Concentration, xMIC
Time, days
Reactor 6
A 0h
B 23h
C 47h
D 76h
E 99h
OD
Concentration, xMIC
Time, days

E

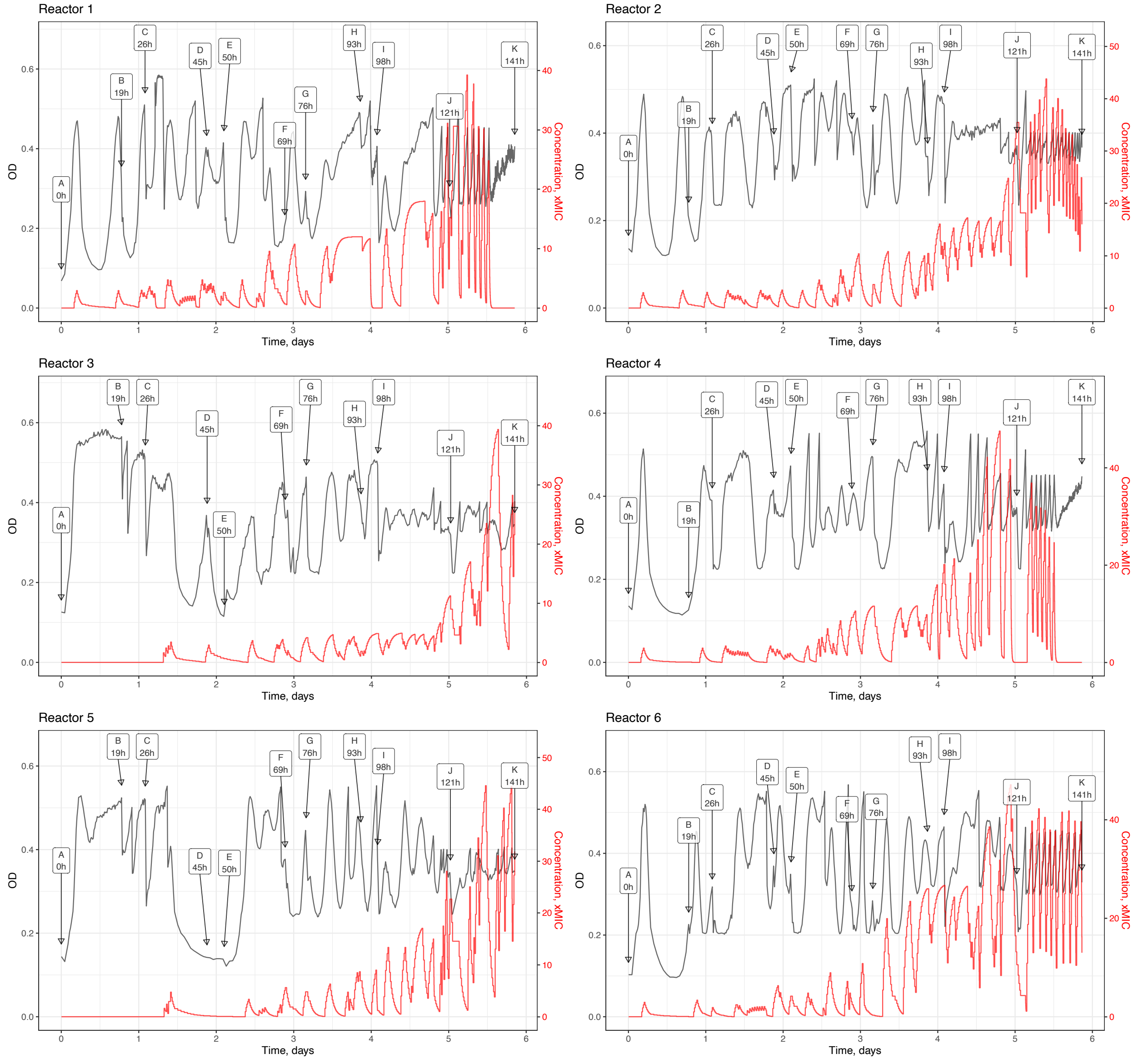
Reactor 1
A 0h
B 19h
C 26h
D 45h
E 50h
F 69h
G 76h
H 93h
I 98h
J 121h
K 141h
OD
Concentration, xMIC
Time, days
Reactor 2
A 0h
B 19h
C 26h
D 45h
E 50h
F 69h
G 76h
H 93h
I 98h
J 121h
K 141h
OD
Concentration, xMIC
Time, days
Reactor 3
A 0h
B 19h
C 26h
D 45h
E 50h
F 69h
G 76h
H 93h
I 98h
J 121h
K 141h
OD
Concentration, xMIC
Time, days
Reactor 4
A 0h
B 19h
C 26h
D 45h
E 50h
F 69h
G 76h
H 93h
I 98h
J 121h
K 141h
OD
Concentration, xMIC
Time, days
Reactor 5
A 0h
B 19h
C 26h
D 45h
E 50h
F 69h
G 76h
H 93h
I 98h
J 121h
K 141h
OD
Concentration, xMIC
Time, days
Reactor 6
A 0h
B 19h
C 26h
D 45h
E 50h
F 69h
G 76h
H 93h
I 98h
J 121h
K 141h
OD
Concentration, xMIC
Time, days
