## Supplementary material for "Shared and Unique Evolutionary Trajectories to Ciprofloxacin Resistance in Gram-negative Bacterial Pathogens": Figure S1

**Supplementary Figure S1. Morbidostat-based experimental evolution workflow.** The initial unevolved culture (1) is plated on the agar Petri dish (2) to make individual colonies. The colonies are collected (3) and used to make 6 inoculates (4) – one colony for each reactor for morbidostat cultivation (5). From each reactor samples are taken roughly once in 24 hours (6). To observe population dynamics (8) we perform WGS with high coverage for each sample (7). The population dynamics is used to choose optimal number of colonies from plated evolved samples (9). For each colony we do WGS (10, 11) and MIC tests (12) that reveals genotype-phenotype association.

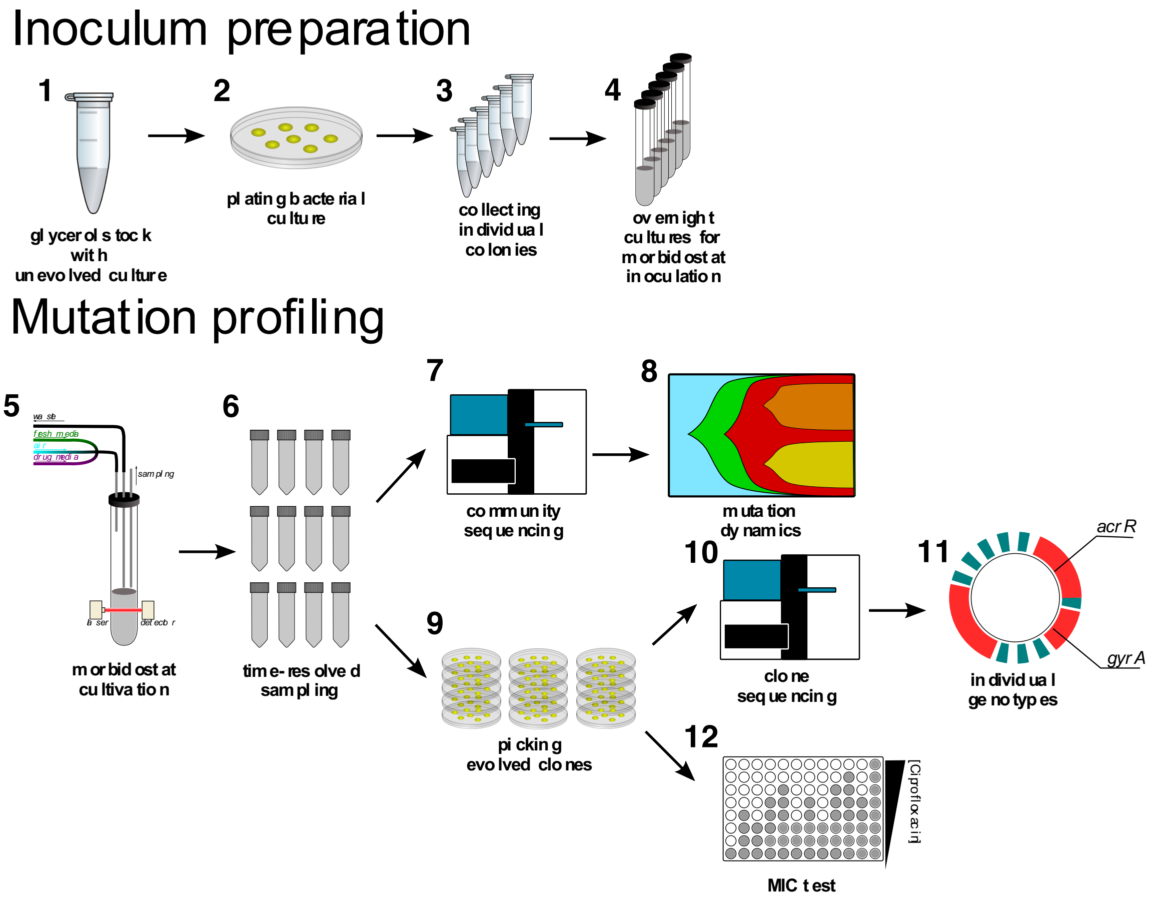
