## Supplementary Methods for "Shared and Unique Evolutionary Trajectories to Ciprofloxacin Resistance in Gram-negative Bacterial Pathogens"

#### I. Morbidostat experiment conditions

##### 1. MORBIDOSTAT RUN CEC-2

**Parental strain:** *Escherichia coli* BW25113.
**MIC for CIP in parental strain:** 0.016 mg/L
**Media without drug (Feed Bottle 1):** MHB
**Media with drug (Feed Bottle 2):** Step 1 (24hrs) – MHB with CIP 0.16 mg/L (10xMIC); Step 2 (44hrs) – MHB with CIP 0.48 mg/L (30xMIC); Step 3 (51hrs) – MHB with CIP 2.4 mg/L (150xMIC).
**Dilution factor:** 20% (4 mL added to 20 mL at each dilution cycle)
**OD thresholds:** LT = 0.15; DT= 0.30
**Dilution cycle:** 15 min at OD>LT; 60 min at OD≤LT
**Sampling:** 10 mL samples taken at time points indicated on the Supplementary Figure S2

##### 2. MORBIDOSTAT RUN CEC-4

**Parental strain:** *Escherichia coli* BW25113.
**MIC for CIP in parental strain:** 0.016 mg/L
**Media without drug (Feed Bottle 1):** MHB, 2% DMSO
**Media with drug (Feed Bottle 2):** Step 1 (47hrs) – MHB/2% DMSO with CIP 0.01 mg/L (0.625xMIC); Step 2 (7hrs) – MHB/2% DMSO with CIP 0.02 mg/L (1.25xMIC); Step 3 (39hrs) – MHB/2% DMSO with CIP 0.08 mg/L (5xMIC); Step 4 (24hrs) – MHB/2% DMSO with CIP 0.32 mg/L (20xMIC); Step 5 (26hrs) – MHB/2% DMSO with CIP 1.28 mg/L (80xMIC);
**Dilution factor:** 20% (4 mL added to 20 mL at each dilution cycle)
**OD thresholds:** LT = 0.15; DT= 0.30
**Dilution cycle:** 15 min at OD>LT; 60 min at OD≤LT
**Sampling:** 10 mL samples taken at time points indicated on the Supplementary Figure S3

##### 3. MORBIDOSTAT RUN CAB-1

**Parental strain:** *Acinetobacter baumannii* ATCC17978.
**MIC for CIP in parental strain:** 0.156 mg/L
**Media without drug (Feed Bottle 1):** MHB, 2% DMSO
**Media with drug (Feed Bottle 2):** Step 1 (7 hrs) – MHB/2% DMSO with CIP 0.195 mg/L (1.25xMIC); Step 2 (43hrs) – MHB/2% DMSO with CIP 0.78 mg/L (5xMIC); Step 3 (20hrs) – MHB/2% DMSO with CIP 1.56 mg/L (10xMIC); Step 4 (23hrs) – MHB/2% DMSO with CIP 6.24 mg/L (40xMIC);
**Dilution factor:** 20% (4 mL added to 20 mL at each dilution cycle)
**OD thresholds:** LT = 0.15; DT= 0.30
**Dilution cycle:** 15 min at OD>LT; 60 min at OD≤LT
**Sampling:** 10 mL samples taken at time points indicated on the Supplementary Figure S4

##### 4. MORBIDOSTAT RUN PAC-1

**Parental strain:** *Pseudomonas aeruginosa* ATCC27853
**MIC for CIP in parental strain:** 0.16 mg/L
**Media without drug (Feed Bottle 1):** MHB
**Media with drug (Feed Bottle 2):** MHB with CIP.

In the PAC-1 run we used a morbidostat modification allowed to set changes of dilution concentrations independently.

| CIP Concentration | | Time of concentration change, hours after the run start | | | | | |
|---|---|---|---|---|---|---|---|
| **mg/L** | **xMIC** | **Reactor 1** | **Reactor 2** | **Reactor 3** | **Reactor 4** | **Reactor 5** | **Reactor 6** |
| 0.35 | 5 | 69 | 67 | 58 | 69 | 67 | 69 |
| 0.525 | 7.5 | 0 | 2 | 11 | - | 2 | - |
| 0.7 | 10 | 7 | 6 | 5 | 5 | 2 | 15 |
| 1.4 | 20 | 4 | 14 | 7 | 11 | 14 | 4 |
| 2.8 | 40 | 3 | 10 | 2 | 4 | 4 | 11 |
| 3.5 | 50 | 16 | - | 16 | 10 | 10 | - |

**Dilution factor:** 20% (4 mL added to 20 mL at each dilution cycle)
**OD thresholds:** LT = 0.15; DT= 0.30
**Dilution cycle:** 15 min at OD>LT; 60 min at OD≤LT
**Sampling:** 10 mL samples taken at time points indicated on the Supplementary Figure S5

##### 5. MORBIDOSTAT RUN PAC-2

**Parental strain:** *Pseudomonas aeruginosa* ATCC27853
**MIC for CIP in parental strain:** 0.16 mg/L
**Media without drug (Feed Bottle 1):** MHB
**Media with drug (Feed Bottle 2):** MHB with CIP.
In the PAC-2 run we used a morbidostat modification allowed to set changes of dilution concentrations independently.

| CIP Concentration | | Time of concentration change, hours after the run start | | | | | |
|---|---|---|---|---|---|---|---|
| **mg/L** | **xMIC** | **Reactor 1** | **Reactor 2** | **Reactor 3** | **Reactor 4** | **Reactor 5** | **Reactor 6** |
| 0.8 | 5 | 29 | 58 | 21 | 56 | 22 | 43 |
| 1.28 | 8 | 34 | 7 | 97 | 3 | 11 | 18 |
| 1.92 | 12 | 33 | 27 | 6 | 34 | 65 | 7 |
| 2.88 | 18 | 20 | 24 | 6 | 3 | 18 | 5 |
| 4.32 | 27 | 2 | 2 | 2 | 10 | 2 | 36 |
| 6.56 | 41 | 2 | 5 | 8 | 3 | 1 | 7 |
| 8 | 50 | 21 | 18 | 1 | 32 | 22 | 25 |

**Dilution factor:** 20% (4 mL added to 20 mL at each dilution cycle)
**OD thresholds:** LT = 0.10; DT= 0.20
**Dilution cycle:** 15 min at OD>LT; 60 min at OD≤LT
**Sampling:** 10 mL samples taken at time points indicated on the Supplementary Figure S6

#### II. Sequencing data analysis

##### 1. Software list

All computations were made on CentOS v.7.6.

- R v. 3.6.0 (https://www.r-project.org/)
- Python 3.6.7 (https://www.python.org/)

- FastQC v. 0.11.8 (https://www.bioinformatics.babraham.ac.uk/projects/fastqc/) [1]
- Trimmomatic v. 0.36 (http://www.usadellab.org/cms/?page=trimmomatic) [2]
- BWA v. 0.7.13 (http://bio-bwa.sourceforge.net/) [3]
- Picard v. 2.2.1 (https://broadinstitute.github.io/picard/)
- LoFreq v. 2.1.3.1 (https://csb5.github.io/lofreq/) [4]
- Samtools + bcftools v. 1.3 (http://www.htslib.org/) [3, 5]
- Genome analysis Toolkit (GATK) v. 3.5 (https://software.broadinstitute.org/gatk/) [6, 7]
- Qualimap v. 2.2 (http://qualimap.bioinfo.cipf.es/) [8]
- snpEff v. 4.3 (http://snpeff.sourceforge.net/) [9]
- breseq v. 0.33.2 (http://barricklab.org/breseq) [10]
- CNOGpro (R package, installation through CRAN) [11]
- Albacore v. 2.3.4 (https://community.nanoporetech.com/; Customer login required)
- Prechop v. 0.2.4 (https://github.com/rrwick/Porechop)
- MinionQC v. 1.3.0 (https://github.com/roblanf/minion_qc) [12]
- SPAdes v. 3.13.0 (http://cab.spbu.ru/software/spades/) [13]
- MUMmer v. 3.1 (http://mummer.sourceforge.net/) [14]
- iJump (https://github.com/sleyn/ijump)

**Custom scripts (available on GitHub):**

- `timeline.py`
- `CNOGpro.R`
- `breseq_parser_confident.py`
- `spread_gd_tsv.R`
- `make_repeats.pl`

#### 2. Reads quality control

We accessed read quality with FastQC software [1]. The Q-scores in all samples had a very good distribution across all positions in reads and we did not trim reads based on Q-scores. The adapters were trimmed with Trimmomatic [2]:

```
java -jar trimmomatic-0.36.jar PE \
    -phred33 \
    [Input reads 1] \
    [Input reads 2] \
    [Trimmed output paired reads 1] \
    [Trimmed output single reads 1] \
    [Trimmed output paired reads 1] \
    [Trimmed output single reads 2] \
    ILLUMINACLIP:adapters.fasta:2:30:10
    MINLEN:65
```

The adapter sequences used for trimming were copied from the Illumina Adapter Sequences document for the corresponding library preparation kit. Additionally, we added overrepresented sequences to the adapters fasta file as we found that it improves read quality. These overrepresented sequences in most cases were adapters with barcodes.

#### 3. Population sequencing data analysis

##### 3.1. Align reads on reference

Processed reads were aligned to a reference genome with BWA-MEM [15]. First, we should create read group header that is required for Genome Analysis Toolkit (GATK) [6]:

```
read1=[trimmed paired reads 1]
rgid=$(echo $read1)
rgid=${rgid/_1_trimmed.fq.gz/}        # Leave only ID
rg="@RG\\tID:$rgid\\tSM:S1\\tPL:illumina\\tLB:L001\\tPU:R1"
```

Then run BWA-MEM itself:

```
bwa mem \
    -t [Number of CPUs] \
    -M \       # Parameter used for Picard compatibility
    -R "$rg" \ # Read group header
    -Y \       # Write all split hits as soft clipped
    [path to the reference fasta] \
    [Trimmed paired reads 1] \
    [Trimmed paired reads 2] > Sample.sam
```

Sort SAM file, save it as BAM and create index with Picard tools (https://broadinstitute.github.io/picard/):

```
java -jar ~/ngsbin/picard-tools-2.2.1/picard.jar SortSam \
    I=Sample.sam \
    O=Sample.bam \
    SO=coordinate \
    CREATE_INDEX=true
```

##### 3.2. Alignment recalibration

Realign reads with LoFreq Vitebri algorithm [4]. As reads positions could be changed, we repeat sorting on the output bam:

```
lofreq viterbi \
    --ref [Path to the reference fasta] \
    --keepflags \
    -o Sample_v.bam \
    Sample.bam
samtools sort -o Sample_i.bam Sample_v.bam
samtools index -b Sample_i.bam
```

We performed the recommended Base Quality Score Recalibration (BQSR) with GATK. However, as there is no database of know SNP for *A. baumannii* and *E. coli* are available. The official advise from GATK is to make a first round of calling first and choose the most trusted mutations. However, as we have a high coverage samples and most trusted mutations would be anyway with high frequency we decided to skip it and make BQSR with the empty SNP database:

```
java -jar ~/ngsbin/GATK-3.5/GenomeAnalysisTK.jar \
    -T BaseRecalibrator \
    -R Path to the reference fasta] \
    -I Sample_i.bam \
    -knownSites known.vcf \
    -o Sample_re.bqsr.grp

java -jar ~/ngsbin/GATK-3.5/GenomeAnalysisTK.jar \
    -T PrintReads \
    -R [Path to the reference fasta] \
    -I Sample_i.bam \
    -BQSR Sample_re.bqsr.grp \
    -o Sample_r_bqsr.bam
```

To assess alignment quality we used Qualimap [8]:

```
qualimap bamqc \
    -bam Sample_r_bqsr.bam \
    -outfile BamQC.pdf
```

##### 3.3. SNP and indel variant calling

The variant calling of SNP and indels was performed with LoFreq:

```
lofreq call-parallel \
    --pp-threads [Number of CPUs] \
    --call-indels \
    -f [Path to the reference fasta] \
    -o variants.lofreq.vcf \
    Sample_r_bqsr.bam
```

##### 3.4. IS elements rearrangement

- The search of mobile elements rearrangement was done with iJump (https://github.com/sleyn/ijump) – the software that we developed to search for IS elements rearrangements in a population sequencing data. In general algorithm of iJump is following:

1. Find soft-clipped reads near boundaries of mobile elements.
2. Extract unaligned part of reads.
3. BLAST unaligned parts against the reference.

4. Find best hits with >90% identity to the reference with unique highest bitscore. If two or more hits share the same bit score - the junction considered as ambiguous and skipped. Usually this is happening because aligner maps reads to several copies of mobile element.
5. Assess frequency of insertion by a formula:

$$Frequency = \frac{\frac{R_l + R_r}{2} * (1 + \frac{B_{min}}{A_{rlen}})}{D_t * (1 - \frac{mmatch}{A_{rlen}})}$$

where:

$R_l$ – a number of reads that support junction to the target on the "left" side of mobile element.

$R_r$ – a number of reads that support junction to the target on the "right" side of mobile element.

*Dt* – average depth of coverage of the target region.

$B_{min}$ – a threshold for the minimum length for unaligned parts of the reads used in the BLAST step. In algorithm we used 10nt.

$A_{rlen}$ – an average read length;

*mmatch* – a minimum length of the read part that could be aligned to reference. The *mmatch* parameter is accessed from the data as a minimum of longest clipped part of the read (e.g. for read in the SAM file with CIGAR string 10S120M30S mmatch is 30).

Before using iJump we made a BLAST search of reference genomes against ISFinder database [16]. The output HTML files were processed with the isfinder_parse.py script that comes with the iJump.

```
python3 isfinder_parse.py \
    -i [ISFinder BLAST HTML page]
```

The script filters BLAST output and creates a table of IS elements boundaries that is used by iJump. The iJump is running with the following command:

```
python3 isjump.py \
    -a [Alignment BAM of SAM file] \
    -r [Reference FNA file with nucleotide FASTA] \
    -g [Reference GFF file with annotations]
    -i [IS coordinates]
```

Detailed description of iJump usage and input data formats could be on the GitHub page (https://github.com/sleyn/ijump).

##### 3.5. Variant annotation and mutational dynamics analysis

To annotate variant calling results we used snpEff [9] and custom scripts. All VCF files from LoFreq output were copied to one directory and merged into one VCF file using bcftools:

```
for file in *.vcf
do
    gz=${file/.vcf/.gz}
    bgzip -c $file > $gz
    tabix -p vcf $gz
done

bcftools merge \
    -m none \
    -o merged.vcf \
    *.gz
```

snpEff was applied on the merged file:

```
java -jar ~/ngsbin/snpEff/snpEff.jar \
    -ud 100 \
    -no-downstream \
    -no-upstream \
    -v [database for the selected reference genome] \
    merged.vcf > ann.vcf
```

Results of the LoFreq variant calling were combined into one comparative table using custom `timeline.py` script.

```
python3 timeline.py \
    -g [Reference GFF file with annotations] \
    -a ann.vcf \
    -r [File with repeats (see below)] \
    -v [Path to the directory with VCF files] \
    -o summary_table.txt
```

Additionally, to manual curation of the variant list we mark two cases of suspicious variants:

1. Variants in the potential repetitive regions where mapped reads could be attributed to the wrong copy of the region by error.

   To do this we followed instructions from MUMmer [14] manual to find repeats in the reference genome:

   ```
   prefix=[Prefix of the output files]

   nucmer \
        -p $prefix \
        -nosimplify \
        --maxmatch \
        [Reference FNA file] \
        [Reference FNA file]
   show-coords \
   ```

```
        -clrT \
        -I 90 \
        -L 65 \
        $prefix".delta" > \
        $prefix".aln"

    make_repeats.pl $prefix
```

This procedure will create file with potential repeated regions in the genome with identity 90% or more and length at least 65nt.

2. Variants that have stable less than 100% frequency in the population. It is another sign that alignment software has troubles with this region.
   To test uniformity of frequencies of a variant across samples for each reactor we preform chi-squared test for uniform distribution. This test id done automatically with the `timeline.py` script.

Results of the iJump IS element rearrangement calling were combined to a comparative table using combine_results.py script from the iJump package.

```
python3 combine_results.py \
    -d [Path to the directory with iJump report files]
```

#### 4. Clonal sequencing data analysis

##### 4.1. Variant calling

We used the breseq software [10] to call variants in clonal data:

```
breseq \
    -j [number of CPUs] \
    -n [Clone ID] \
    -r [Reference GBK file] \
    --require-match-length 40 \
    [Trimmed paired reads 1] \
    [Trimmed paired reads 2]
```

The Copy Number Variation analysis was done using CNOGpro package for the R programming language. The CNOGpro package uses GBK files but it is limited for using files with one contig only. To use CNOGpro it is necessary to split multiple contigs GBK file to the separate files for each contig. For each contig the hitsfile containing coordiantes of mapped reads should be generated (for details see CNOGpro manual):

```
# in breseq data directory for each contig:
export CONTIG=[Contig name]
samtools view reference.bam | \
    perl -lane \
```

```
            'print "$F[2]\t$F[3]" \
            if $F[2] =~ m/$ENV{CONTIG}/' \
        > out.hits
```

CNOGpro was wrapped by us into an R script. Script filters out IS elements and outputs only CDS:

```
  Rscript --vanilla CNOGpro.R \
      [Reference GBK file with the selected contig] \
      [Contig name] \
      [iJump IS elements coordinates table to filter out
them] \
      [Clone ID used for output file name]
```

##### 4.2. Mutational data comparison

To collect breseq results in the table format we used custom script that makes TSV table from the annotated GD files ([breseq work directory]/output/evidence/annotated.gd):

```
  python3 breseq_parser_confident.py \
      -g [directory with annotated gd files] \
      -o [output file]
```

To make a comparison table for variants found in the clonal sequencing data we used gdtools with a custom script for reformatting the gdtools output:

```
  gdtools ANNOTATE \
      -r [Reference GBK file] \
      -o [Output file] \
      -f TSV \                # TSV format
      [Directory with GD files]/*.gd

  Rscript --vanilla spread_gd_tsv.R \
      [gdtools output file] \
      [Output table]
```

To make a comparison table for copy number variations found with CNOGpro we used a custom script:

```
  Rscript --vanilla Reshape_cnv.R \
      [Directory with the CNOGpro outputs] \
      [Output table] \
      [Minimum difference from 1 to show gene in the output.
We used 0.3]
```

#### 5. Novel assembly of *Acenitobacter baumannii* ATCC 17978 genome

Initial variant calling against public reference revealed high number of variants in unevolved culture of *Acinetobacter baumannii* ATCC 19978 (number of variants with frequency >85%: 16 variants against GCA_001593425.2, 98.46% mapped reads; 87 variants against GCA_000015425.1, 98.97% mapped reads). To improve variant calling we decided to make a hybrid assembly with Oxford Nanopore long reads.

One clone of unevolved *Acinetobacter baumannii* ATCC 19978 was sequences with ONT MinION sequencer using Nanopore Rapid Barcoding gDNA Sequencing kit (SQK-RBK004). Raw sequencing data QC was performed with MinionQC.

Reads were demultiplexed with Albacore basecaller (ONT community site):

```
read_fast5_basecaller.py \
    -i [Path to the FAST5 files directory] \
    -t [Number of CPUs] \
    -s [Path to the directory for output FASTQ file] \
    -f FLO-MIN106 \
    -k SQK-RBK004 \
    -r \
    -o fastq \
    -q 0
```

As in the library all barcoded samples were from the same organism to increase coverage, we decided to make second round of demultiplexing on unclassified reads with Porechop (https://github.com/rrwick/Porechop):

```
porechop \
    -i [Path to the unclassified/ folder] \
    -b Porechop_barcodes/ \
    --threads [Number of CPUs] \
    --untrimmed \
    --discard_middle
```

Adapters from the reads demultiplexed by Albacore were trimmed by Porechop:

```
porechop \
    -i [FASTQ file] \
    -o [Output trimmed FASTQ file] \
    --discard_middle \
    --threads [Number of CPUs]
```

Both reads demultiplexed by Albacore and Porechop were combined by simple cat command. The hybrid assembly of *Acinetobacter baumannii* ATCC 19978 genome was performed using SPAdes assembler [13]:

```
spades.py \
    -k 21,33,55,77 \
```

```
--careful \
--nanopore [Nanopore fasta files] \
[Trimmed Illumina paired reads 1] \
[Trimmed Illumina paired reads 2]
```

Assembled genome was annotated by RASTtk pipeline on the corresponding web server (https://rast.nmpdr.org/) [17].

### References


1. Andrews S. FastQC. A quality control tool for high throughput sequence data.). Babraham Institute (2010).

2. Bolger AM, Lohse M, Usadel B. Trimmomatic: a flexible trimmer for Illumina sequence data. *Bioinformatics* **30**, 2114-2120 (2014).

3. Li H*, et al.* The Sequence Alignment/Map format and SAMtools. *Bioinformatics* **25**, 2078-2079 (2009).

4. Wilm A*, et al.* LoFreq: a sequence-quality aware, ultra-sensitive variant caller for uncovering cell-population heterogeneity from high-throughput sequencing datasets. *Nucleic Acids Res* **40**, 11189-11201 (2012).

5. Li H. A statistical framework for SNP calling, mutation discovery, association mapping and population genetical parameter estimation from sequencing data. *Bioinformatics* **27**, 2987-2993 (2011).

6. DePristo MA*, et al.* A framework for variation discovery and genotyping using next-generation DNA sequencing data. *Nat Genet* **43**, 491-498 (2011).

7. Van der Auwera GA*, et al.* From FastQ data to high confidence variant calls: the Genome Analysis Toolkit best practices pipeline. *Curr Protoc Bioinformatics* **43**, 11 10 11-11 10 33 (2013).

8. Okonechnikov K, Conesa A, Garcia-Alcalde F. Qualimap 2: advanced multi-sample quality control for high-throughput sequencing data. *Bioinformatics* **32**, 292-294 (2016).

9. Cingolani P*, et al.* A program for annotating and predicting the effects of single nucleotide polymorphisms, SnpEff: SNPs in the genome of Drosophila melanogaster strain w1118; iso-2; iso-3. *Fly (Austin)* **6**, 80-92 (2012).

10. Deatherage DE, Barrick JE. Identification of mutations in laboratory-evolved microbes from next-generation sequencing data using breseq. *Methods Mol Biol* **1151**, 165-188 (2014).

11. Brynildsrud O, Snipen LG, Bohlin J. CNOGpro: detection and quantification of CNVs in prokaryotic whole-genome sequencing data. *Bioinformatics* **31**, 1708-1715 (2015).

12. Lanfear R, Schalamun M, Kainer D, Wang W, Schwessinger B. MinIONQC: fast and simple quality control for MinION sequencing data. *Bioinformatics* **35**, 523-525 (2019).

13. Bankevich A*, et al.* SPAdes: a new genome assembly algorithm and its applications to single-cell sequencing. *J Comput Biol* **19**, 455-477 (2012).

14. Kurtz S*, et al.* Versatile and open software for comparing large genomes. *Genome Biol* **5**, R12 (2004).

15. Li H, Durbin R. Fast and accurate short read alignment with Burrows-Wheeler transform. *Bioinformatics* **25**, 1754-1760 (2009).

16. Siguier P, Perochon J, Lestrade L, Mahillon J, Chandler M. ISfinder: the reference centre for bacterial insertion sequences. *Nucleic Acids Res* **34**, D32-36 (2006).

17. Brettin T*, et al.* RASTtk: a modular and extensible implementation of the RAST algorithm for building custom annotation pipelines and annotating batches of genomes. *Sci Rep* **5**, 8365 (2015).
